# STEER-seq: a recombinase based platform for pooled multimodal single-cell perturbation screening

**DOI:** 10.64898/2026.10.08.757551

**Authors:** Valentina Migliori, Chenqu Suo, Olli Dufva, Ivan Gyulev, Krzysztof Polanski, Vasiliki Symeonidou, Geraldine Rodschinka, Ioannis Sarropoulos, Sho Iwama, Alexander S. Steemers, Aleksander M. Gontarczyk, Hayden Powell, Ana-Maria Cujba, Jong-Eun Park, Mathew J. Garnett, George S. Vassiliou, Sarah A. Teichmann, Andrew R. Bassett

**Author notes:** These authors contributed equally (Valentina Migliori, Chenqu Suo, Olli Dufva). These authors contributed equally (Sarah A. Teichmann, Andrew R. Bassett). Corresponding authors: Valentina Migliori, Sarah A. Teichmann and Andrew Bassett.

## Abstract

Pooled gain-of-function screening is a powerful method for screening for transcription factors whose overexpression leads to changes in cellular identity. However, variable transgene copy number, integration site and genomic context from random viral integration can confound quantitative comparisons between perturbations. We present STEER-seq (Single-cell Transgene Expression via Exchange Recombination sequencing), a virus-free platform for pooled ORF gain-of-function screening in human induced pluripotent stem cells (hiPSCs). STEER-seq combines irreversible Bxb1-mediated integration at a defined genomic safe-harbour locus with promoterless donor libraries and barcode-enabled perturbation tracking, enabling single-copy expression from a common promoter and genomic context and direct recovery of perturbation identity by single-cell sequencing. In a focused haematopoietic screen, SPI1 expression alone was sufficient to drive differentiation towards HSC/MPP-like progenitors in the absence of exogenous cytokines, with transient induction generating progenitors that retained multilineage haematopoietic differentiation potential. Joint RNA-chromatin accessibility profiling linked transcription factor-induced transcriptional responses to chromatin remodelling and enabled reconstruction of perturbation-induced gene regulatory networks, including a RUNX2-driven osteoblast programme. Scaling STEER-seq to around 2,000 human coding sequences enabled systematic discovery of transcription factors capable of inducing distinct cell-state programmes, including neural progenitor-, melanocyte- and vascular smooth muscle-like states, as well as maintenance of pluripotency. STEER-seq provides a scalable, quantitative framework for single-cell and multimodal gain-of-function screening, enabling functional discovery and engineering of human cell identity.

## Introduction

The ability to systematically identify genes capable of controlling cellular identity is central to functional genomics, disease modelling and cell engineering. Transcription factors (TFs) are particularly powerful perturbations because they regulate coordinated gene-expression programmes and can be sufficient to drive major changes in cell state. A striking example is somatic cell reprogramming, where forced expression of only four TFs (OCT4, SOX2, KLF4 and MYC) can override an established somatic transcriptional programme and generate human induced pluripotent stem cells (hiPSCs)^1^. Importantly, such transitions do not always require combinations of factors: overexpression of a single lineage-specifying TF, such as NGN2, is sufficient to rapidly drive human iPSCs towards a neuronal fate^2^. These observations provide a strong rationale for systematic screening of TFs to identify those capable of directing cells towards desired states and to generate important training data for modelling and prediction of factors that drive cell fate transitions.

Large-scale gain-of-function screens have demonstrated the potential of this approach. Lentiviral open reading frame (ORF) libraries enable thousands of TFs to be screened in parallel, including in hiPSCs and with single-cell transcriptomic readouts^3,4^. More recently, high-multiplicity lentiviral delivery has extended this approach to combinatorial TF screening, enabling exploration of large TF-combination landscapes for cell-state engineering, including haematopoietic progenitor-like states^5^. However, lentiviral delivery does not control transgene copy number or genomic integration site generating variability in expression level and number of TFs per cell^6–10^. These variables are particularly important when comparing the capacity of different TFs to drive cell-state changes or for comparison between defined variant ORFs. Expressing each ORF as a single copy from the same promoter and defined genomic context reduces variation arising from transgene dosage and integration site, allowing phenotypic differences to be more directly attributed to the TF or variant being tested. Viral approaches additionally require library packaging, production and transduction which introduce biases against longer ORFs due to limits on packaging capacity in lentivirus, adding complexity to both pooled screening and downstream validation. A virus-free approach operating directly from plasmid libraries therefore offers practical as well as quantitative advantages.

Exogenous TF overexpression also addresses a conceptually distinct question from CRISPR activation (CRISPRa), which activates genes from their endogenous loci and whose efficiency depends on guide RNA activity, promoter architecture and chromatin accessibility^11–14^. For cell-state engineering, the aim is not necessarily to reproduce physiological TF expression or native developmental trajectories, but to identify factors whose expression is sufficient to drive cells towards a desired state. The capacity of a TF to induce a particular cell state cannot necessarily be inferred from its endogenous expression alone, and unbiased gain-of-function screening can reveal unexpected TF–cell-state relationships^4^. Exogenous ORF screening additionally enables defined splice isoforms^15^ and coding variants to be tested independently of endogenous promoter regulation.

Site-specific recombinase-based landing pads provide a route to achieve controlled, virus-free library integration. The serine recombinase Bxb1 enables efficient and irreversible integration of large DNA payloads^16–20^. The scalability of this approach was demonstrated through site-specific, single-copy integration of sgRNA libraries in CHO cells for pooled CRISPR loss-of-function screening^21^. Bxb1 landing pads have also been employed in HEK293T cells for functional analysis of protein-variant libraries and their combinations^21,22,23^, and this has been further developed using orthogonal Bxb1 sites and double landing pads for protein-variant assays^23,24^. In parallel, Bxb1-based systems have been established at predefined genomic loci in hiPSCs, including the CLYBL safe-harbour locus and platforms such as STRAIGHT-IN^17,25^ and Big-IN^26^, primarily for precise genome engineering and generation of isogenic cell lines. However, to date it has not been possible to perform large-scale gain-of-function screening in hiPSCs with single cell molecular readouts, which is important to understand cell-state transitions in these pluripotent cells. Existing large-scale TF gain-of-function screens in hiPSCs rely on viral delivery, and have achieved scale and single-cell transcriptomic phenotyping, but do not control transgene copy number and genomic context^3–5^.

Here we present STEER-seq (Single-cell Transgene Expression via Exchange Recombination sequencing), a virus-free pooled gain-of-function screening platform that combines Bxb1-mediated integration at the CLYBL safe-harbour locus with barcoded ORF libraries and single-cell molecular readouts in hiPSCs. STEER-seq expresses perturbations from a common promoter and predefined genomic context while enabling their identities to be recovered directly by single-cell RNA sequencing. Extension of perturbation capture to joint RNA and chromatin-accessibility profiling further enables the transcriptional and regulatory consequences of TF overexpression to be measured simultaneously, linking perturbations to changes in cis-regulatory elements and gene regulatory networks. Single- and dual-landing-pad configurations additionally enable controlled individual and pairwise perturbations from defined genomic contexts, providing a complementary strategy to high-MOI combinatorial screening^5^.

We applied STEER-seq across targeted, multimodal and large-scale gain-of-function screens during hiPSC differentiation. In a focused haematopoietic screen, we found that SPI1 expression was sufficient to drive differentiation towards HSC/MPP-like progenitors in the absence of exogenous cytokines. Transient SPI1 induction generated progenitors that retained multilineage haematopoietic differentiation potential following withdrawal of the instructive signal. Extending STEER-seq to joint RNA and chromatin accessibility profiling enabled simultaneous measurement of the transcriptional and regulatory consequences of TF overexpression. For example, RUNX2 induced an osteoblast-associated transcriptional programme together with extensive changes in chromatin accessibility, enabling reconstruction of perturbation-induced enhancer-driven gene regulatory networks. Finally, scaling STEER-seq to more than 2,000 human TFs enabled systematic discovery of factors driving distinct cell identities, including SMAD7-induced neural progenitor-like states, NR4A3-associated melanocyte-like differentiation and NR4A2-induced vascular smooth muscle-like states, alongside factors promoting maintenance of pluripotency. Together, these results establish STEER-seq as a quantitative, virus-free framework for single-cell and multimodal gain-of-function screening, functional discovery and engineering of human cell identity.

## Results

### Development of STEER-seq for quantitative pooled gain-of-function screening with single cell readout

To establish a quantitative, virus-free gain-of-function screening platform in hiPSCs, we first targeted a Bxb1-mediated landing-pad to the permissive *CLYBL* safe-harbour locus using CRISPR-mediated homology-directed repair (Fig. 1a). The cassette contained a constitutive CAG promoter driving Bxb1 expression as well as drug and fluorescent selectable markers, flanked by attP recombination sites. Correct integration at the *CLYBL* locus was confirmed by targeted long read sequencing (Extended Data Fig. 1a, b). Donor constructs were designed without a promoter and contained an attB-flanked transgene, puromycin-resistance cassette and transcribed perturbation barcode (Fig. 1b). Following Bxb1-mediated cassette exchange, the resident landing-pad cassette is replaced by the donor, positioning the transgene downstream of the genomic CAG promoter. Thus, transgene expression is activated following genomic integration while expression from non-integrated plasmid DNA is minimised. Because Bxb1 recombination converts attP and attB into attL and attR sites that can no longer undergo recombination, each landing pad can undergo a single irreversible integration event^18,27^. We isolated hiPSC clones carrying either one or two landing-pad copies, corresponding to monoallelic and biallelic *CLYBL* targeting, respectively, and confirmed copy number by droplet digital PCR (ddPCR) (Fig. 1c). To assess cassette-exchange efficiency, we introduced a promoterless GFP donor into single-landing-pad cells. Following puromycin selection, 94.2% of cells were GFP-positive, demonstrating efficient donor integration and transgene expression from the genomic landing pad (Fig. 1d). Targeted long-read sequencing across the *CLYBL* locus further confirmed the expected cassette-exchange structure at both recombination junctions (Extended Data Fig. 1c).To assess expression stability, we monitored constitutive mApple expression from the landing pad during differentiation. Expression initially decreased, but subsequently recovered and remained readily detectable until week 7 (Extended Data Fig. 1d), indicating that the locus remains transcriptionally active during differentiation.

**Figure 1.**
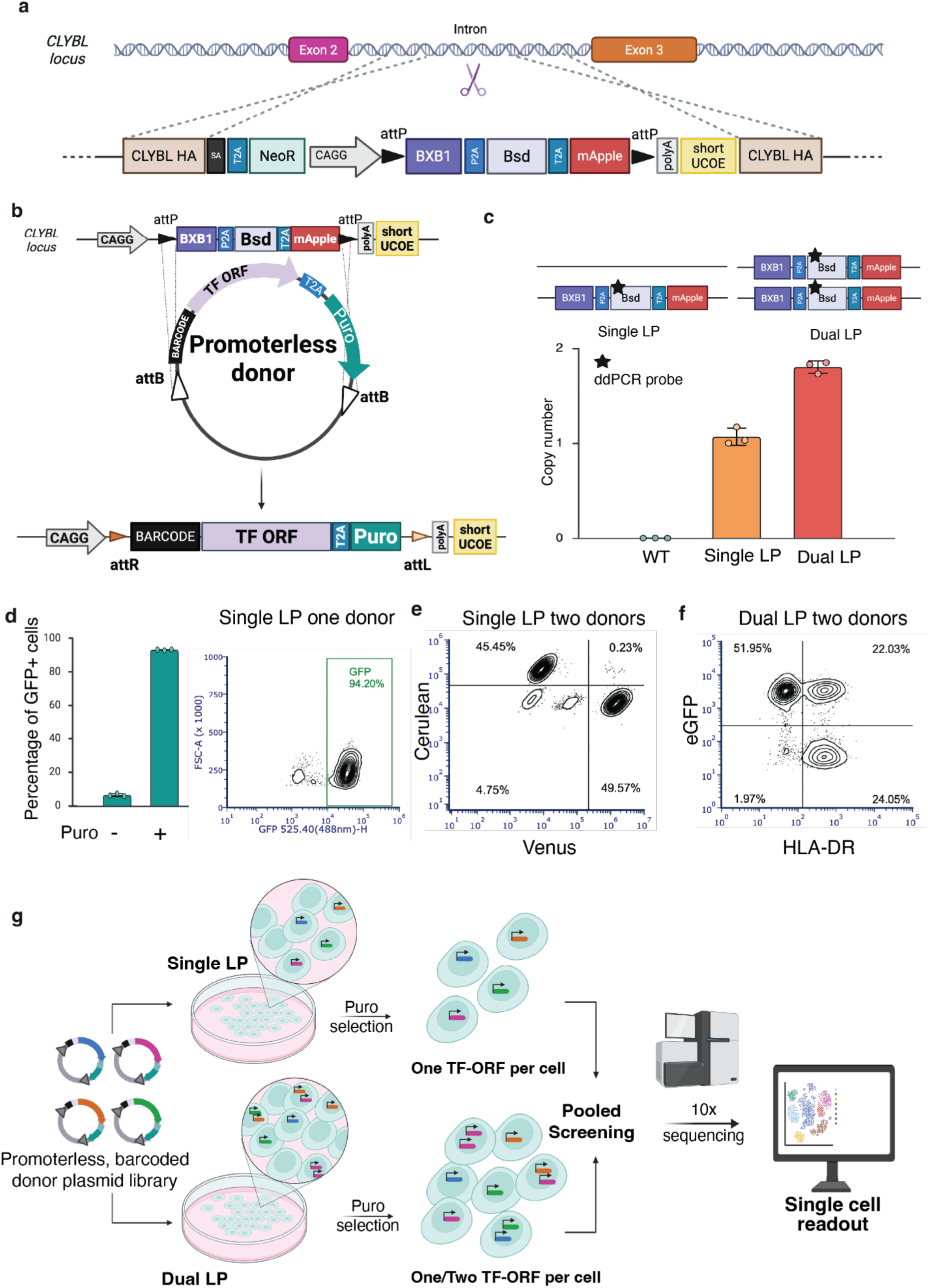
Method overview. **a,** Schematic of the Bxb1 recombinase-compatible landing pad integrated into the human *CLYBL* safe-harbour locus. The landing pad contains a constitutively expressed Bxb1-P2A-Blasticidin-T2A-mApple cassette flanked by attP recombination sites and is targeted to the intronic region between exons 2 and 3. HA, homology arm; SA, splice acceptor; NeoR, neomycin resistance; Bsd, blasticidin resistance; UCOE, ubiquitous chromatin opening element. **b,** Schematic of Bxb1-mediated donor integration. Promoterless donor constructs containing a transcription factor (TF) payload, puromycin resistance cassette and barcode sequence are flanked by attB recombination sites. Following unidirectional cassette exchange, the donor is integrated into the genomic landing pad downstream of the resident CAG promoter, enabling expression only after correct genomic integration. **c,** Determination of landing pad copy number by droplet digital PCR (ddPCR). Representative single landing-pad (single LP) and dual landing-pad (dual LP) configurations are shown above. ddPCR analysis identified clones carrying either one or two landing pad integrations corresponding to monoallelic and biallelic targeting events, respectively. Points represent biological replicates and bars indicate mean ± s.d. **d,** Assessment of cassette exchange efficiency in single landing-pad cells using a promoterless GFP donor. Following Bxb1-mediated integration and puromycin selection, GFP expression was quantified by flow cytometry. Left, percentage of GFP-positive cells with or without puromycin selection. Right, representative flow cytometry plot showing GFP-positive cells following cassette exchange. Bars indicate mean ± s.d.; points represent biological replicates. **e,** Validation of one-gene-per-cell integration in the single landing-pad line. Cells carrying a single landing pad were co-transfected with two promoterless donor plasmids encoding Venus or Cerulean together with Bxb1 recombinase. Flow cytometry shows mutually exclusive Venus- or Cerulean-positive populations, demonstrating that only a single donor cassette can integrate into the single landing pad despite simultaneous delivery of multiple donors. **f,** Validation of dual independent integrations in the dual landing-pad line. Cells carrying two landing pads were co-transfected with GFP and CIITA promoterless donor constructs. Flow cytometry for GFP expression and HLA-DR antibody staining identifies GFP-only, CIITA-only and GFP/CIITA double-positive populations, demonstrating independent occupancy of the two landing pads and enabling simultaneous expression of two defined transgenes. **g,** Experimental workflow for pooled transcription factor screening using single and dual landing pad cell lines. Barcoded promoterless donor libraries are introduced into landing pad cells by Bxb1-mediated recombination and selected with puromycin. Single landing-pad cells acquire one integrated perturbation per cell, whereas dual landing-pad cells can acquire one or two perturbations. Perturbation identity is recovered by single-cell sequencing for downstream analysis. Images in panels a, b and c were created in BioRender. Bassett, A. (2026) https://BioRender.com/zom28mh

We next tested whether the single-landing-pad configuration restricts integration to one perturbation per cell. Single-landing-pad cells were co-transfected with promoterless Cerulean and Venus donors. Cells expressed either Cerulean or Venus, but not both, consistent with mutually exclusive occupancy of the single landing pad despite simultaneous exposure to multiple donor constructs (Fig. 1e). Conversely, co-transfection of dual-landing-pad cells with GFP and CIITA donors generated GFP-only, HLA-DR-only and GFP/HLA-DR double-positive populations. As CIITA is the master transcriptional regulator of MHC class II genes, successful CIITA expression induces cell-surface HLA-DR, enabling HLA-DR staining to serve as a phenotypic readout of CIITA activity. The presence of all three populations therefore demonstrated the capacity of dual-landing-pad cells to acquire and express two distinct perturbations, supporting the feasibility of controlled pairwise perturbation screening from defined genomic contexts (Fig. 1f). The total fraction of cells undergoing dual integration is likely greater than the 22% observed as double-positive by FACS, as cells receiving the same donor at both landing-pad alleles cannot be distinguished by this assay.

Thus, cells carrying a single landing pad are configured to express one transgene, whereas dual-landing-pad cells can express up to two independently introduced perturbations, providing an architecture for defined single and pairwise gain-of-function studies.

Finally, to link each integrated ORF directly to its single-cell phenotype, donor constructs contain a transcribed perturbation barcode recoverable by single-cell sequencing. We developed barcode architectures compatible with both 5′ and 3′ capture, enabling perturbation assignment from single-cell transcriptomic and multiomic readouts such as RNA and chromatin accessibility^28,29^(Fig. 1g). STEER-seq therefore combines virus-free pooled ORF delivery, controlled integration from a predefined genomic context and direct single-cell perturbation assignment. Single-landing-pad cells restrict integration to one perturbation per cell, while dual-landing-pad cells can support up to two perturbations from the same pooled library, enabling controlled pairwise gain-of-function screening. This provides a complementary strategy to high-MOI viral approaches that maximise combinatorial diversity by introducing multiple TFs per cell. Because successful cassette exchange replaces the mApple-expressing landing-pad cassette, loss of mApple fluorescence can additionally be used to enrich dual-landing-pad cells in which both landing-pad cassettes have been disrupted by recombination.

### Targeted transcription factor screening identifies *SPI1* as a potent inducer of haematopoietic differentiation in the absence of other external signals

To evaluate the utility of the landing pad platform for pooled gain-of-function screening during differentiation, we performed a targeted transcription factor (TF) screen using a previously established protocol^30^ for the generation of haematopoietic progenitors from hiPSCs (Fig. 2a). We selected 12 candidate TFs that are highly expressed in *in vivo* haemogenic endothelial and haematopoietic populations^31,32^, generated a pooled donor library together with an *EGFP* control and transfected our iPSC line containing a single landing-pad. Following Bxb1-mediated integration and puromycin selection, cells were differentiated under either standard cytokine-supported conditions or cytokine-free conditions to assess the ability of individual TFs to promote haematopoietic specification. Samples were collected at D0, D8 and D18 and profiled by single-cell RNA sequencing.

**Figure 2.**
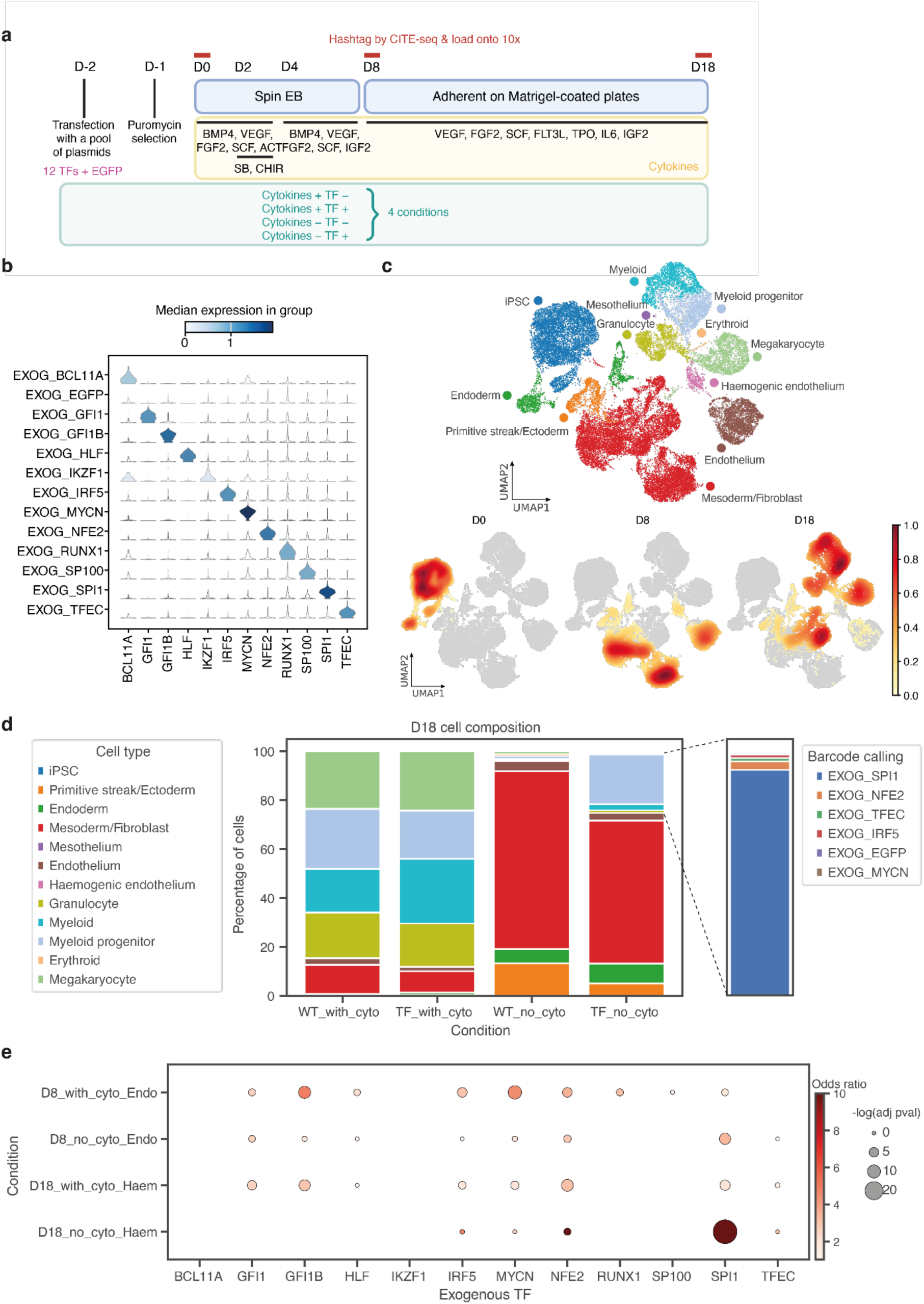
Targeted HSC screen reveals *SPI1* as promising target. **a,** Experimental design for targeted HSC screening. Twelve TFs enriched in *in vivo* haemogenic endothelium or HSCs, together with EGFP, were pooled at equal ratios and transfected into iPSCs on day −2 (D−2). Transfected cells underwent 24 h puromycin selection from D−1 to D0 before differentiation. Cells were differentiated following the protocol of Ng et al.^30^ through spin embryoid body (EB) formation and subsequent transfer onto Matrigel-coated plates at D8. Cells were cultured either in the presence or absence of haematopoietic cytokines, generating four conditions: wild type (WT) with cytokines (cytokine+TF−), TF pool with cytokines (cytokine+TF+), WT without cytokines (cytokine−TF−), and TF pool without cytokines (cytokine+TF+). Cells were harvested at D0, D8 and D18, with samples from different conditions multiplexed using hashtag antibodies prior to loading onto the 10x Genomics platform for single-cell RNA sequencing. **b,** Stacked violin plots showing log-normalised expression of each TF across cells assigned to the corresponding exogenous TF barcode. **c,** UMAP embedding of integrated scRNA-seq datasets collected across all differentiation conditions and sampling time points, coloured by annotated cell types (top). Bottom panels show density projections of cells from individual time points overlaid onto the integrated UMAP, illustrating the temporal progression of differentiation trajectories. **d,** Stacked bar plots showing the proportion of annotated cell types in each differentiation condition at D18. The rightmost bar shows exogenous barcode assignments among haematopoietic-lineage cells from the TF+/cyto− condition, highlighting enrichment of *SPI1*-overexpressing cells within the haematopoietic compartment. **e,** Dot plot showing enrichment test results for exogenous TF barcode representation across selected differentiation conditions and cell types. Dot colour represents the odds ratio of enrichment, whereas dot size corresponds to −log(adjusted *P*-value). Dot sizes are shown only for TF-cell type pairs with adjusted *P*-values ≤ 0.05 and odds ratios > 1. For visualisation purposes, odds ratios greater than 10 were capped at 10.

Perturbation identity was assigned using integrated barcode sequences captured during single-cell RNA sequencing. The majority of cells (97.3%) with detectable perturbation barcodes only contained a single barcode (Extended Data Fig. 2a) showing the precision of STEER-seq in generating single copy integrations. For each cell, exogenous TF identity was assigned based on the barcode with the highest UMI count, and requiring the dominant barcode to account for more than 50% of all detected perturbation barcode counts. Cells assigned to each perturbation showed selective overexpression of the corresponding exogenous TF, confirming accurate perturbation assignment and robust recovery of perturbation identity (Fig. 2b). STEER-seq therefore enables perturbation assignment directly from single-cell transcriptome libraries without additional enrichment, even with modest barcode transcript abundance (Extended Data Fig. 2b).

Across all conditions and time points, we obtained 35,017 high-quality single-cell transcriptomes spanning multiple stages of mesodermal and haematopoietic differentiation. Unsupervised clustering identified populations corresponding to iPSCs, primitive streak/ectoderm, endoderm, mesoderm/fibroblast, mesothelium, endothelium, haemogenic endothelium and downstream haematopoietic lineages, including erythroid, megakaryocyte, granulocyte, myeloid progenitor and myeloid populations (Fig. 2c and Extended Data Fig. 2c and d). Cells treated with cytokines followed the expected developmental progression from pluripotent and mesodermal states at early time points towards endothelial and haematopoietic populations at later stages of differentiation.

We next examined the effects of cytokine supplementation and TF overexpression on lineage output. As expected, cytokine-supported cultures efficiently generated haematopoietic progenitor and mature blood cell populations by D18 (Fig. 2d). In contrast, cytokine-free cultures were predominantly composed of mesodermal/fibroblast-like and endothelial populations, indicating a failure to efficiently progress towards haematopoietic fates. Notably, TF-overexpressing cells regained the capacity to generate downstream haematopoietic populations. This effect was most evident for *SPI1* and, to a lesser extent, *NFE2*, which promoted the emergence of haematopoietic cell types even in the absence of exogenous cytokine stimulation (Fig. 2d).

To systematically quantify TF-specific lineage biases, we performed enrichment analysis comparing the representation of each perturbation within individual cell types relative to the *EGFP* control. Several TFs showed modest enrichment within endothelial or haematopoietic compartments; however, *SPI1* exhibited the strongest and most consistent effect across differentiation stages and culture conditions (Fig. 2e). *SPI1*-overexpressing cells were significantly enriched within endothelial populations at D8 and haematopoietic populations at D18 under both cytokine-supported and cytokine-free conditions. The strongest enrichment was observed within D18 haematopoietic populations generated in the absence of cytokines, indicating that *SPI1* can promote haematopoietic differentiation even when exogenous developmental cues are removed. Together, these results identify *SPI1* as the most potent inducer of haematopoietic differentiation among the TFs tested and nominate it for further functional investigation.

To assess the viability of the dual landing-pad system in a screening context, we performed an additional experiment using dual landing-pad iPSCs transfected with a library containing five TFs and *EGFP*. Barcode recovery identified 3,767 cells carrying a single unique barcode and 520 cells carrying two distinct barcodes (Extended Data Fig. 2e). Cells assigned to individual or paired barcodes showed concordant overexpression of the corresponding transgenes, confirming accurate recovery of perturbation identity. These results demonstrate that the dual-landing-pad system can generate cells with robust overexpression of defined exogenous TF combinations, supporting its future use for controlled pairwise gain-of-function screening in which the effects of individual TFs can be compared directly with the corresponding TF combinations.

### Transient *SPI1* induction generates functional HSC/MPP-like progenitors in the absence of cytokines

To validate TFs identified in the pooled screen, we selected five of the strongest candidates associated with endothelial and haematopoietic populations (GFI1, GFI1B, HLF, NFE2 and SPI1) and repeated the screen in an arrayed format using constitutive expression. Following 18 days of differentiation, cells were enriched for haematopoietic stem and progenitor cells (HSPCs, Lin−CD34+CD90+, Extended Data Fig. 3a)^33^, and subjected to single-cell RNA sequencing (Extended Data Fig. 3b).

Automated cell type annotation using CellTypist^34^ identified populations corresponding to HSC/multipotent progenitor (HSC/MPP) and cycling MPP states (Extended Data Fig. 3c). These populations expressed canonical HSC/MPP-associated genes, including *CD34*, *SPINK2*, *HOPX* and *RUNX1* (Extended Data Fig. 3d). Consistent with the pooled screen, SPI1 generated the highest proportion of HSC/MPP-like cells among the five candidates tested (Extended Data Fig. 3e). SPI1/PU.1 is an established regulator of haematopoietic specification and HSPC maintenance and a key regulator of myeloid lineage specification, and has previously been implicated in hiPSC-derived haematopoiesis, including as part of TF combinations promoting HSPC generation^35–37^. Thus, although the identification of SPI1 itself was not unexpected, our results demonstrate that SPI1 alone, rather than as part of a multi-TF combination, and in the absence of exogenous cytokine stimulation can strongly promote an HSC/MPP-like transcriptional state.

Because sustained *SPI1* expression has been associated with myeloid lineage commitment^38–40^, we reasoned that transient SPI1 activation might be sufficient to establish an HSC/MPP-like progenitor state while its subsequent withdrawal could permit broader lineage differentiation. We therefore generated a doxycycline-inducible SPI1 line using the landing-pad platform (Fig. 3a), in which Bxb1-mediated cassette exchange placed SPI1 expression under the control of a doxycycline-responsive promoter. Efficient donor integration was confirmed following puromycin selection, yielding >95% GFP-positive cells (Fig. 3b), and doxycycline-dependent expression was validated by flow cytometry (Fig. 3c), reaching substantially higher levels than constitutive expression.

**Figure 3.**
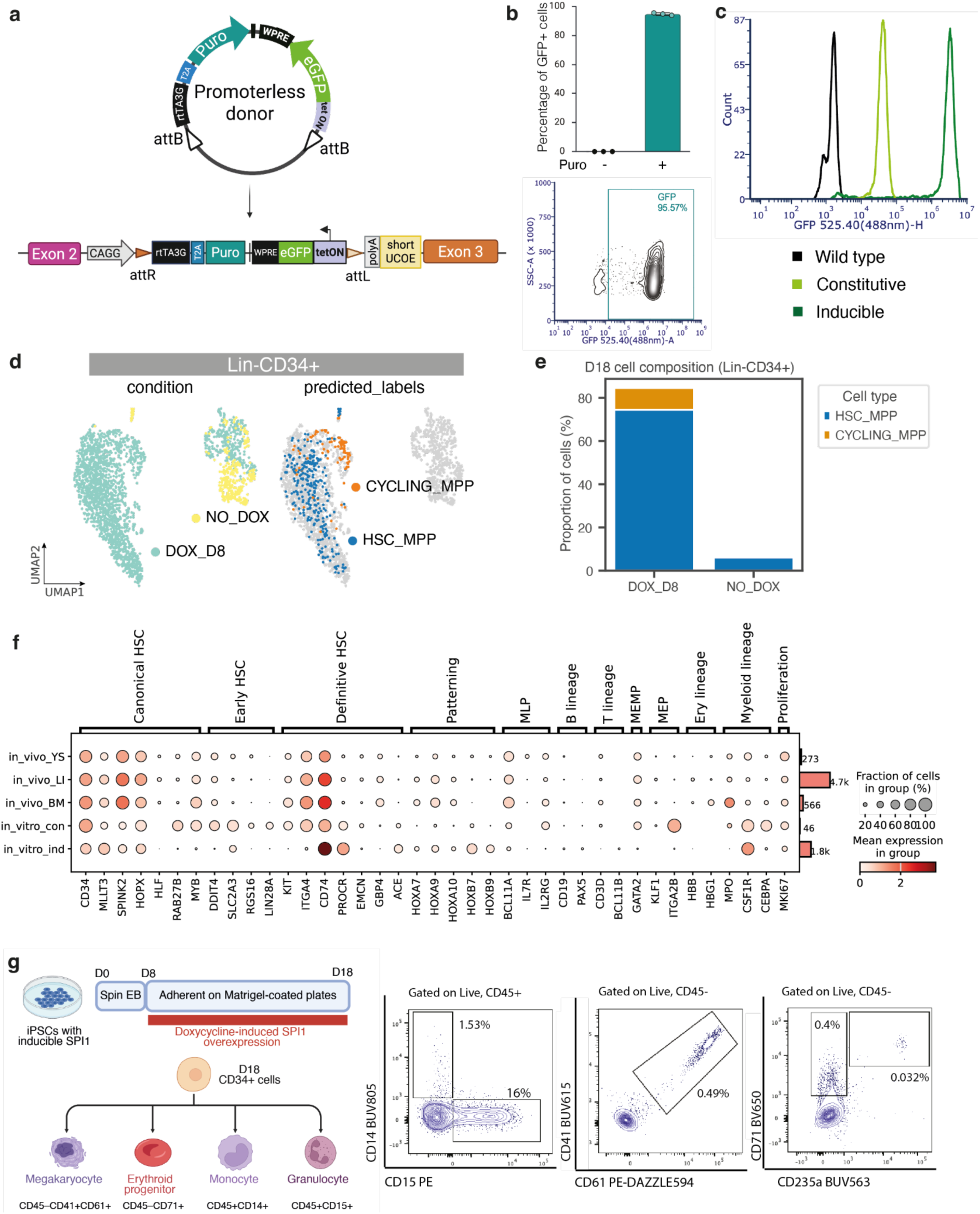
Validation of *SPI1* using inducible promoter. **a,** Schematic of the inducible overexpression system in the landing-pad hiPSC line. Site-specific recombination mediates single-copy integration of a doxycycline-inducible GFP reporter, enabling controlled induction and tracking of expressing cells by fluorescence. Upon doxycycline treatment, the integrated GFP is expressed from the targeted genomic locus. Created in BioRender. Bassett, A. (2026) https://BioRender.com/zom28mh **b,** Validation of donor integration and selection efficiency. Top, percentage of GFP-positive cells before and after puromycin selection and induction with 2 µg/mL doxycycline for 24h, demonstrating enrichment of correctly integrated cells. Bottom, representative flow-cytometry analysis of GFP expression following selection.**c,** Flow-cytometry analysis of GFP expression in inducible and constitutive overexpression lines. Representative fluorescence intensity histograms show robust transgene expression in both configurations, with inducible expression activated by with 2 µg/mL doxycycline treatment. **d,** UMAP embedding of scRNA-seq data from sorted Lin−CD34+ cells harvested at D18 from inducible *SPI1* cultures with with 2 µg/mL doxycycline treatment between D8 and D18 (DOX_D8) or untreated controls (NO_DOX). Cells are coloured by differentiation condition (left) and CellTypist-predicted cell type using the Pan Fetal Human model^43^ (right). Only cells predicted as HSC_MPP or CYCLING_MPP are highlighted in the cell type panel; all other cells are shown in grey. **e,** Stacked bar plots showing the proportion of manually annotated HSC_MPP and CYCLING_MPP cells among D18 Lin−CD34+ cells in the DOX_D8 and NO_DOX conditions. The remaining cells, not shown in the bar plots, are predominantly endothelial cells. Cell type annotations were assigned based on the leiden clustering and marker gene expression shown in Extended Data Fig. 4b-d. **f,** Dot plot showing the expression of selected marker genes across *in vitro* and *in vivo* HSC/MPP populations, with *in vivo* reference data obtained from Suo et al. (2022)^43^. Dot colour indicates mean gene expression (log-normalised counts), and dot size represents the fraction of cells expressing each gene. Rows correspond to cell populations and columns correspond to marker genes. The total number of cells in each cell population is shown in the accompanying bar plot on the right. Marker genes were selected based on previous studies by Suo et al. (2022)^43^ and Goh et al. (2023)^55^. In_vitro_con comprises cells from constitutive *SPI1* overexpression cultures predicted as HSC_MPP or CYCLING_MPP by CellTypist majority voting. In_vitro_ind comprises cells from inducible *SPI1* overexpression cultures treated with with 2 µg/mL doxycycline between D8 and D18 and manually annotated as HSC_MPP or CYCLING_MPP (Extended Data Fig. 4d). *In vivo* populations comprise HSC_MPP and CYCLING_MPP cells from yolk sac (YS), fetal liver (LI), and bone marrow (BM) samples. **g,** Functional assessment of haematopoietic progenitor potential generated by inducible SPI1 overexpression. D18 CD34+ cells, purified by magnetic cell separation (MACS), derived from inducible *SPI1* cultures treated with with 2 µg/mL doxycycline from D8 to D18 were subjected to lineage differentiation assays. Left, experimental schematic and lineage outputs. Right, flow cytometry plots showing generation of monocytic, granulocytic, megakaryocytic, and erythroid populations, identified by the indicated surface marker combinations. Gating was performed as described in Extended Data Fig. 4e. The percentages on the plots indicate the frequencies of the total live cell population.

Based on previous studies implicating *SPI1* during early haematopoietic specification^41,42^, doxycycline induction was initiated at day 8 of differentiation and maintained until day 18 under cytokine-free conditions. Cells were subsequently enriched for a hematopoietic population (Lin−CD34+, Extended Data Fig. 4a) and profiled by single-cell RNA sequencing. Analysis revealed a marked shift in cell identity following *SPI1* induction. Label transfer using CellTypist classified the majority of induced cells as HSC/MPP-like or cycling MPP populations, whereas cells from the non-induced control condition were predominantly annotated as endothelial cells (Fig. 3d and e, Extended Data Fig. 4b–d). Importantly, in the absence of SPI1, very few cells progressed into the hematopoietic progenitor compartment (Lin−CD34+). These findings indicate that transient *SPI1* activation following establishment of endothelial intermediates is sufficient to drive progression towards haematopoietic progenitor states without exogenous cytokine stimulation.

To further assess the molecular identity of these cells, we compared *SPI1*-induced HSC/MPP-like cells to HSC-enriched populations from human fetal cross-tissue atlas^43^ (Fig. 3f). *SPI1*-induced cells expressed multiple genes associated with human HSCs and multipotent progenitors, including *CD34*, *MLLT3* and *HOPX*, together with definitive HSC markers such as *CD74* and *PROCR*. Although the transcriptional profiles did not fully recapitulate those of fetal HSCs, with lower expression of *SPINK2* and limited expression of *RAB27B* and *MYB*, the emergence of HSC/MPP-like populations under cytokine-free conditions indicates that *SPI1* alone is sufficient to activate a substantial component of the haematopoietic stem and progenitor cell transcriptional programme. In addition, *SPI1*-induced progenitors also expressed the myeloid-associated receptor *CSF1R*, consistent with early myeloid priming and the established role of SPI1 in myeloid specification.

We next asked whether cells induced into this progenitor state retained differentiation potential following withdrawal of SPI1 expression. As sustained SPI1 expression can promote myeloid commitment, withdrawal allowed us to test whether the induced progenitor state could persist independently of continued ectopic SPI1 expression while retaining the capacity to respond to lineage-specific differentiation cues. Doxycycline was removed at day 18 and CD34+ cells were cultured under lineage-specific differentiation conditions (Fig. 3g) that were capable of supporting hematopoietic differentiation of cord-blood-derived CD34+ cells (14.6% myeloid, 2.7% erythroid, 9.6% granulocytic). SPI1-induced progenitors generated cells expressing markers of megakaryocytic, erythroid and myeloid lineages, including monocyte and granulocyte populations (Fig. 3g and Extended Data Fig. 4e). As expected from the role of SPI1 in myeloid specification, differentiation was biased towards myeloid output, but erythroid and megakaryocytic populations were still detected albeit at lower frequencies. Thus, transient SPI1 expression was sufficient to establish an HSC/MPP-like progenitor state under cytokine-free conditions, while subsequent withdrawal of SPI1 revealed detectable differentiation potential across multiple haematopoietic lineages.

Together, these results independently validate SPI1 as the strongest haematopoietic regulator identified in the pooled screen and demonstrate that temporally restricted expression of a single TF can promote an HSC/MPP-like state under cytokine-free conditions that retains differentiation potential following withdrawal of the inducing TF.

### Multi-omic compatibility enables simultaneous screening of transcriptional and chromatin perturbations

We next aimed to broaden the compatibility of our approach to other readouts such as single-nucleus multiomics measuring gene expression and chromatin accessibility from the same cells (10x Multiome). The combined RNA and ATAC multiomic readout is particularly suitable for our TF-based screening, as it can identify factors that alter chromatin accessibility and enable inference of enhancer-linked gene regulatory networks downstream of TF induction.

We engineered a version of the donor vector where an additional barcode was incorporated at the 3’ end close to the polyadenylation signal, allowing barcode detection in 3’-based assays such as 10x Multiome, and many other single cell multiomic methodologies such as Parse Biosciences, Illumina, etc. (Fig. 4a).

**Figure 4.**
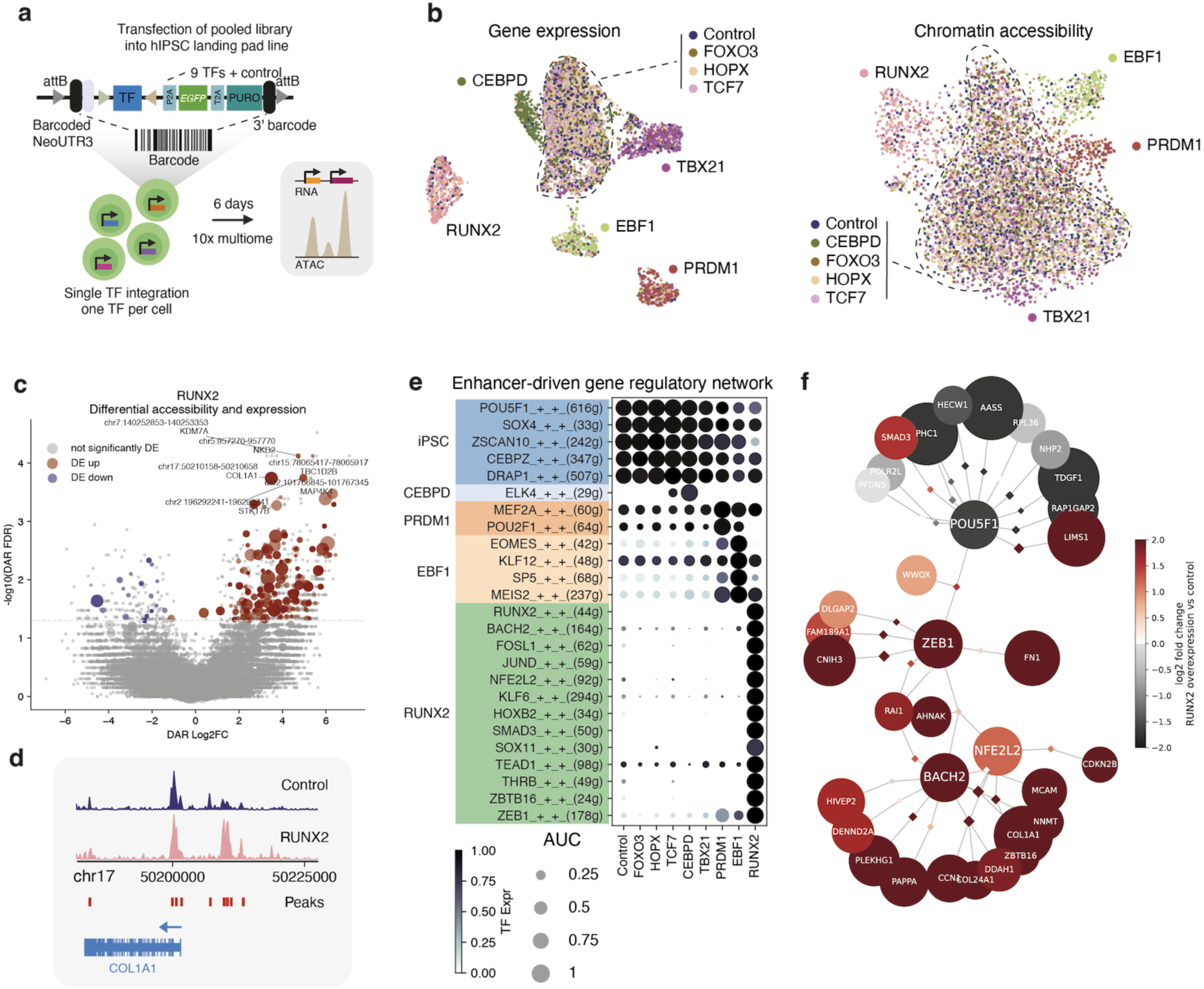
TF screening with joint gene expression and chromatin accessibility readout. **a,** Schematic of the vector harbouring barcodes both 5’ and 3’ relative to the TF, and the pooled overexpression experiment in iPSC with single-nucleus RNA/ATAC multiome readout. Created in BioRender. Bassett, A. (2026) https://BioRender.com/zom28mh **b,** UMAP embeddings based on gene expression and chromatin accessibility in iPSCs with pooled overexpression of 9 TFs and control (6 days post-transfection), with overexpressed TF based on detected barcodes coloured on the individual nuclei. **c,** Differentially accessible regions between nuclei overexpressing RUNX2 and controls, with differential accessibility log2 fold change on x axis and significance on y axis. Significance threshold at 0.05 FDR is indicated with a dashed line. Regions where the linked gene (based on SCENIC+ region to gene links) is significantly (FDR<0.05) differentially expressed in a concordant fashion are coloured red or blue, with higher dot size indicating higher absolute value of the differential expression log2 fold change. Most significantly differentially accessible regions with linked genes are labelled. **d,** Chromatin accessibility tracks at a genomic region surrounding the *COL1A1* gene encoding type I collagen, an osteoblast marker and highly differentially expressed and accessible gene between control and RUNX2-overexpressing nuclei. Peaks called by ArchR are shown as vertical red lines, and the location of the gene is shown at the bottom with exons marked as vertical lines and direction of transcription indicated by an arrow. **e,** Dot plot showing selected SCENIC+-derived regulons out of the most highly variable ones across perturbations, grouped by perturbation. Dot size indicates regulon activity (AUCell score based on gene expression) and colour intensity indicates transcription factor expression. Number of genes belonging to each regulon is indicated next to the TF. Regulons are grouped and coloured by the overexpressed TF most strongly inducing the regulons. **f,** Network plot showing a subset of TF-region-target links impacted by RUNX2 overexpression. Most highly variably expressed target genes (indicated in small font size) and highly variably accessible regions (diamonds) for selected TFs (indicated in large font size) induced (*BACH2*, *NFE2L2*, *ZEB1*) or downregulated (*POU5F1*) by RUNX2 overexpression are shown. Colour indicates differential expression (circles) or accessibility (diamonds) log2 fold change between RUNX2-overexpressing and control cells, and larger size indicates higher significance.

To evaluate the feasibility and the opportunities of the multiome readout, we cloned a pool of 9 TFs into the 3’ barcoded vector, along with a Venus fluorescent protein as a control, and transfected the pool into iPSCs carrying a single landing-pad. After 6 days, including 3 days of puromycin selection, we isolated nuclei and performed multiomic profiling. We obtained 9,942 nuclei with both high-quality RNA and ATAC data, 6,192 of which had a detectable barcode, indicating direct applicability of our technology for the multiome readout (Extended Data Fig. 5a).

Assigning TF perturbations to nuclei based on the highest barcode count in each cell, we observed clustering of cells carrying 5 TFs (*RUNX2*, *EBF1*, *PRDM1*, *TBX21*, *CEBPD*) into distinct clusters based on gene expression (Fig. 4b). For chromatin accessibility, we called peaks using ArchR based on the TF perturbation groups and similarly evaluated clusters in the ATAC latent semantic indexing-based space. We found three TFs (*RUNX2*, *EBF1*, *PRDM1*) inducing distinct clusters in the ATAC space, indicating that these TFs may harbour pioneering capacity for changing the chromatin landscape, in contrast to *TBX21* and *CEBPD* which may act predominantly by binding to already open chromatin regions.

We then evaluated differentially accessible regions and differentially expressed genes in perturbed nuclei compared to control (Extended Data Fig. 5b-d). *RUNX2* overexpression induced the most differential chromatin regions and differentially expressed genes, with many but not all of them overlapping. Thus, capturing both modalities simultaneously provides complementary information about TF-driven gene regulation (Fig. 4c). To evaluate which cell types each TF might direct iPSCs towards, we scored cells in a fetal cross-tissue atlas^43^ using signatures derived from the differentially expressed genes. The *RUNX2*-induced score was highest in osteoblasts (Extended Data Fig. 5e-f), indicating ability of *RUNX2* to drive osteoblast-related gene programmes from iPSC, consistent with increased accessibility of a cis-regulatory element (CRE) near the osteoblast hallmark gene *COL1A1* encoding type I collagen (Fig. 4d). This indicates the utility of our multiome TF screening approach to decipher detailed gene regulatory mechanisms of TFs-mediated gene induction at the level of individual CREs.

The ability to simultaneously measure induced genes and CREs following TF overexpression suggested that our technology could enable construction of perturbation-induced enhancer-driven gene regulatory networks (eGRNs). To achieve this, we inferred eGRNs using SCENIC+ by considering each perturbation as a distinct cell state (Fig. 4e). The resulting eGRN allowed us to identify regulons driven by TFs activated downstream of *RUNX2* overexpression. Changes in chromatin accessibility in regions linked with the FOSL1 and NFE2L2 transcription factors were observed, consistent with increased RNA expression. The analysis also illustrated how *COL1A1* is jointly regulated by *BACH2*, *FOSL1* and *NFE2L2* at the same enhancer, while *ZEB1* drives another osteoblast hallmark *FN1* encoding fibronectin, all with increased accessibility and expression in response to *RUNX2* (Fig. 4f). In summary, the applicability of our technology across modalities such as RNA+ATAC multiome unlocks detailed investigation of enhancer-driven gene regulation by TFs.

### Genome-scale pooled TF-ORF screening enables discovery of candidate cell-fate regulators

Having established compatibility of the landing pad platform with both transcriptomic and multiomic readouts, we next sought to evaluate its scalability for large-scale pooled overexpression screening. To this end, we generated a pooled TF-ORF library comprising 2,089 human coding sequences^44,45^, each linked to a barcode. This was achieved through pooled Gateway cloning into a complex barcoded donor vector and using long-read PacBio sequencing to couple barcodes to transcription factor identity (Fig. 5a). The resulting promoterless donor library was introduced into single landing-pad iPSCs by Bxb1-mediated integration, followed by puromycin selection. Cells were then subjected to the same cytokine-free differentiation protocol described above, with embryoid body formation initiated immediately following selection and differentiation continued for 18 days without addition of lineage-specifying cytokines. Cells were collected immediately following selection (D0) and after 18 days of differentiation (D18) and profiled by single-cell RNA sequencing.

**Figure 5.**
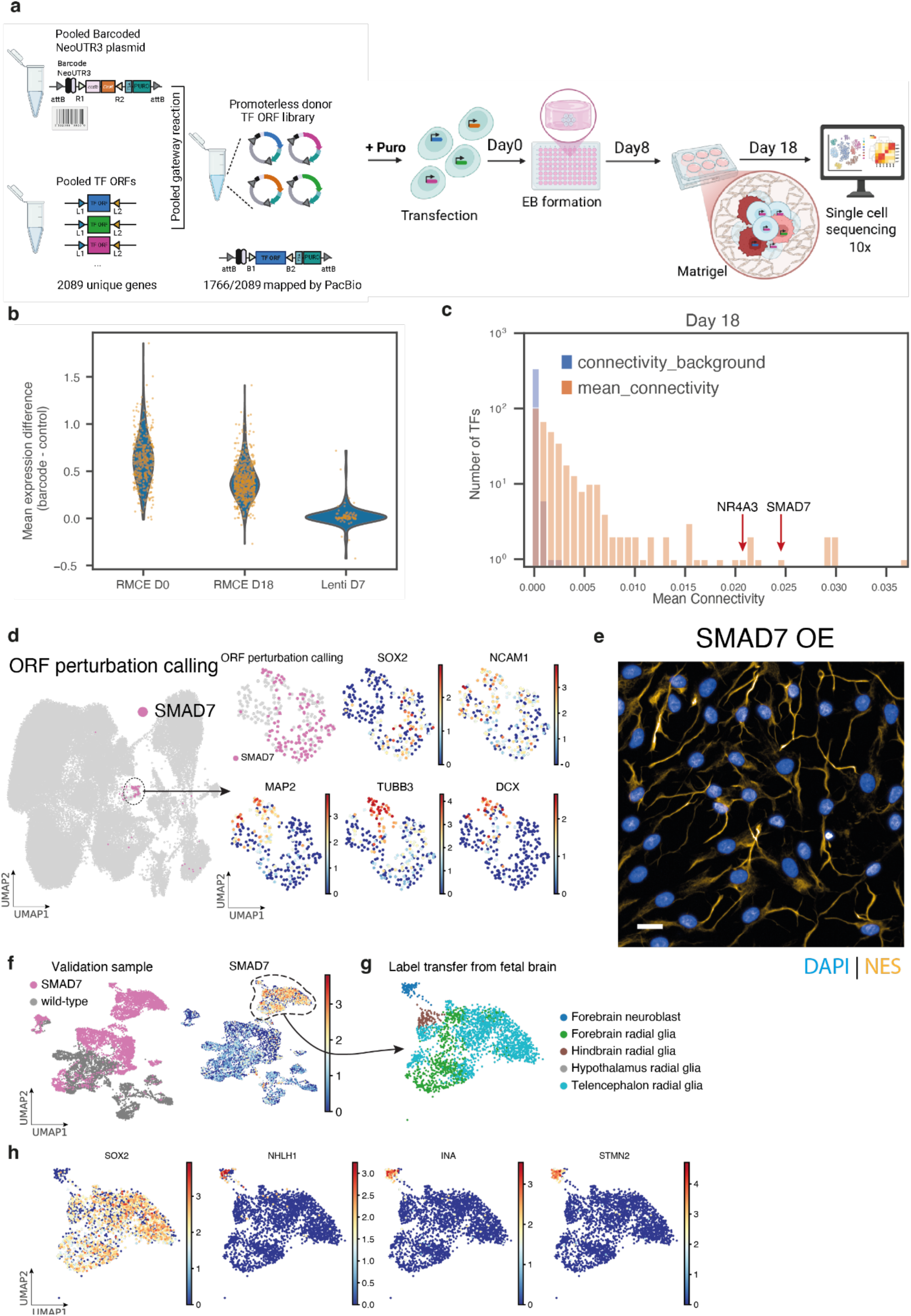
Genome-scale pooled TF-ORF screening and validation of *SMAD7*. **a,** Overview of the pooled ORF overexpression screening workflow. A library of 2,089 human TF open reading frames (ORFs) was cloned into a 5’ barcoded NeoUTR3 donor backbone through a pooled Gateway reaction, generating a pooled barcoded TF-ORF library. Long-read PacBio sequencing was used to map barcodes to TF ORFs, yielding 1,766 uniquely assigned ORF–barcode pairs. Donor plasmid pool was transfected into single landing-pad iPSCs, resulting in site-specific single-copy integration of one ORF construct per cell. Following puromycin selection, cells were aggregated to form embryoid bodies (EBs; day 0), plated onto Matrigel, and differentiated in APEL medium without cytokines. Cells were harvested on day 18 for single-cell RNA-sequencing, enabling assignment of ORF identity through barcode detection and characterisation of TF-induced cell-state transitions. Created in BioRender. Bassett, A. (2026) https://BioRender.com/zom28mh **b,** Violin plots show the distribution of mean log1p-normalised expression differences between cells assigned to each ORF barcode and control cells, defined as cells with no barcode assigned. Only ORFs represented by at least 10 barcode-positive cells were included. RMCE D0 and RMCE D18 represent day 0 and day 18, respectively, of the pooled ORF screening experiment shown in panel **a**. Lenti D7 represents day 7 of differentiation in the lentiviral-based platform using data from Joung et al. 2024^4^. Individual points overlaid on the violin plots represent individual ORFs. For each ORF, the mean expression difference was calculated as the mean log1p-normalised expression of the corresponding ORF in barcode-positive cells minus its mean log1p-normalised expression in control cells. **c,** Histogram showing the mean pairwise connectivity between cells assigned to the same ORF at Day 18 compared with background connectivity. For each ORF represented by at least 20 cells, the mean within-ORF connectivity was calculated from the scVI-based neighbourhood graph. Background connectivity was calculated by randomly sampling the same number of cells from the full Day 18 dataset. The *y*-axis shows the number of ORFs on a log scale. The mean pairwise connectivities for *SMAD7* and *NR4A3* are highlighted. **d,** UMAP embedding of all D18 cells from the pooled screen, highlighting cells assigned to *SMAD7*-associated barcodes (left). Cells from Leiden cluster 22 (Extended Data Fig. 7a) were then subsetted, and the neighbourhood graph and UMAP embedding were recomputed. The resulting subcluster UMAPs (right) are coloured by *SMAD7* barcode assignment or expression of neuronal marker genes. **e,** Immunostaining image of *SMAD7*-overexpressing cells harvested at Day 18 stained for DAPI and NES (scale bar: 20 µm). **f,** UMAP embedding of cells harvested at D18 from wild-type and *SMAD7*-overexpression cultures, coloured by condition (left) and *SMAD7* expression level (right). **g,** UMAP embedding of *SMAD7*-overexpressing cells subsetted from Leiden clusters 0, 5 and 13 in Extended Data Fig. 7e, coloured by CellTypist-predicted cell types using the Developing Human Brain model^48^. **h,** UMAP embedding of *SMAD7*-overexpressing cells, shown on the same embedding as in **g** and coloured by expression of selected neuronal marker genes.

To assign perturbations, barcode expression was quantified directly from single-cell transcriptomes. To account for barcode contamination from ambient cell-free RNA in the suspension (“soup”), barcode-calling thresholds were determined using barcode counts detected in non-cell droplets (Extended Data Fig. 6a). Across the four samples, the 99.9th percentile of non-zero barcode counts in non-cell droplets ranged from 4 to 5 UMIs (see Methods for details), and we therefore defined positive barcode assignment as >5 barcode UMIs. At this threshold, the estimated false-positive rate in non-cell droplets ranged from 3.27 × 10^-6^ to 1.12 × 10^-5^. Using these stringent criteria, 47.6% of all cells could be assigned at least one perturbation, comprising 89.7% of cells at D0 and 41.2% of cells at D18, while 93.4% of assigned cells contained only a single detectable barcode (Extended Data Fig. 6b). Data from all samples were integrated using scVI^46^ (Extended Data Fig. 6c).

To assess the fidelity of perturbation assignment, we compared expression of each ORF between cells assigned to its corresponding barcode and all other cells. At both D0 and D18, the majority of ORFs exhibited higher expression within their assigned cell populations, with most points falling above the diagonal in Extended Data Fig. 6d. This confirmed concordance between barcode assignment and the corresponding gene overexpression, although transgene expression levels were generally lower at D18 than at D0. Consistently, cells called with ORF barcodes were enriched for expression of the matched ORF at both time points (Extended Data Fig. 6e). Ranking each matched ORF against all other ORFs within each ORF barcode group showed enrichment at low rank percentiles, indicating that matched ORFs were often among the most highly expressed ORFs in their assigned cells (Extended Data Fig. 6f). This pattern was broader at D18, consistent with reduced or more heterogeneous transgene expression after differentiation. We further compared the magnitude of ORF overexpression achieved with our recombinase-mediated cassette exchange (RMCE)-based platform with that reported for the lentiviral-based platform of Joung et al. 2023^4^ (Fig. 5b). ORF overexpression was substantially higher with the RMCE platform, including at D18, where expression remained higher than that observed at D7 of differentiation in the Joung et al. dataset.

We next asked whether overexpression of individual ORFs induced coherent changes in cell state. We reasoned that perturbations capable of driving differentiation or cell-state transitions would cause cells carrying the same perturbation to converge towards similar transcriptional profiles. To quantify this effect, we calculated the mean pairwise connectivity between cells assigned to each perturbation using the integrated neighbourhood graph and compared these values to background distributions generated by random sampling of the same number of cells. Across all perturbations, observed connectivities were significantly greater than expected by chance (Wilcoxon rank-sum test, P = 9.63 × 10^-17^; Fig. 5c), indicating widespread transcriptomic convergence of cells upon the same perturbation.

Several perturbations exhibited particularly strong convergence and formed highly distinct transcriptional populations. Among these, *SMAD7* showed one of the highest connectivity scores and generated a discrete cell cluster containing 84% of all cells assigned to *SMAD7*-associated barcodes (Fig. 5d and Leiden cluster 22 in Extended Data Fig. 7a). Examination of this cluster revealed expression of neural ectoderm-associated genes including *SOX2* and *NCAM1*, together with a subset of cells expressing neuronal markers such as *MAP2*, *TUBB3* and *DCX* (Fig. 5d). Consistent with these observations, automated cell type annotation using CellTypist^34^ identified neuronal cell identities within this population (Extended Data Fig. 7b). A subset of cells within this cluster carried barcodes classified as MULTIHIT (Extended Data Fig. 7c) due to ambiguous barcode–ORF assignments in the PacBio reference library. However, detailed examination of these barcodes revealed that they were overwhelmingly associated with *SMAD7*, indicating that the cluster predominantly represented a *SMAD7*-driven transcriptional state. We therefore selected *SMAD7* for independent validation.

To validate the screening result, we repeated the differentiation using *SMAD7* alone. Immunofluorescence staining for Nestin (NES) revealed pronounced neurite-like projections and neural-like network formation in *SMAD7*-expressing cells (Fig. 5e). Comparison with matched control cells confirmed that this phenotype was specifically enhanced following SMAD7 overexpression, despite the presence of NES-positive cells in both conditions (Extended Data Fig. 7d). NGN2 overexpression was included as a positive control and produced a similarly pronounced neural-like morphology (Extended Data Fig. 7d). We further profiled independently generated *SMAD7*-overexpressing cells using 10x FLEX single-cell sequencing. Validation cells with high *SMAD7* expression (Fig. 5f, Leiden clusters 0, 5, 13 in Extended Data Fig. 7e) expressed neural markers including *NES, SOX2*, *MAP2* and *NCAM1* (Extended Data Fig. 7f). Further supporting this neural identity, most cells in this cluster were predicted by SCimilarity^47^ to be glial cells or radial glial cells, consistent with a neural progenitor-like state (Extended Data Fig. 7g).

To further characterise these neural-lineage cells, we performed label transfer using a fetal brain atlas^48^. Most cells were predicted to be radial glia, consistent with a progenitor identity, whereas a subcluster of cells were predicted to be neuroblasts, indicating progression towards neuronal differentiation^48^ (Fig. 5g). These annotations were supported by *SOX2* expression in predicted radial glia, and *NHLH1*, *INA* and *STMN2* expression in predicted neuroblasts (Fig. 5h). Most radial glia were assigned to telencephalon or forebrain identities, consistent with previous reports that *SMAD7* can induce hESCs towards an anterior neural fate^49^. However, a distinct subcluster was predicted to be hindbrain radial glia (Extended Data Fig. 7h) and expressed hindbrain-associated markers, including *PAX3* and *FST* (Extended Data Fig. 7i). This regional heterogeneity was further reflected by mutually exclusive *PAX3* and *PAX6* expression patterns (Extended Data Fig. 7j).

NR4A3 was another transgene with a high connectivity score (Fig. 5c). Cells assigned to NR4A3-associated barcodes showed expression of melanocyte-associated marker genes, including *TYRP1* and *MITF* (Extended Data Fig. 8a and b). In an independent NR4A3-overexpression validation experiment, we identified a subcluster of cells predicted to be melanocytes, which was absent from the wild-type culture (Extended Data Fig. 8c). These cells expressed highly specific melanocyte markers, including *PMEL*, *TYR*, *TYRP1* and *MLANA* (Extended Data Fig. 8d).

Together, these results demonstrate that the landing pad platform supports genome-scale pooled ORF screening with single-cell resolution and enables identification of perturbations that drive coherent cell-state transitions. The discovery and validation of SMAD7 and NR4A3 provide an example of how transcriptomic convergence analysis can be used to prioritise candidate cell-fate regulators from large-scale pooled gain-of-function screens.

### Correlation-based analysis reveals shared perturbation programmes and NR4A2-induced vascular smooth muscle-like differentiation

The large-scale screening dataset generated above also enabled systematic comparison of transcriptional programmes induced by different perturbations. We reasoned that genes driving related biological processes or cell-fate transitions should produce similar transcriptional outputs, even when overexpressed independently. To identify such relationships, we calculated average gene expression profiles for each perturbation across all assigned cells and computed pairwise Pearson correlations between perturbation-level expression signatures, following a strategy similar to that used by Joung et al. 2023^4^. Hierarchical clustering of the resulting correlation matrix revealed multiple groups of perturbations associated with shared transcriptional states (Fig. 6a). One such group contained SOX2, SCN3B and SMARCC2 (Group 2; Fig. 6a and b). Cells assigned to these perturbations localised predominantly within the same transcriptional cluster and exhibited elevated expression of pluripotency-associated genes including *POU5F1* (OCT4), *SOX2* and *NANOG* (Fig. 6b). These findings suggest that overexpression of these factors promotes retention of, or delayed exit from, pluripotent cell states during differentiation.

**Figure 6.**
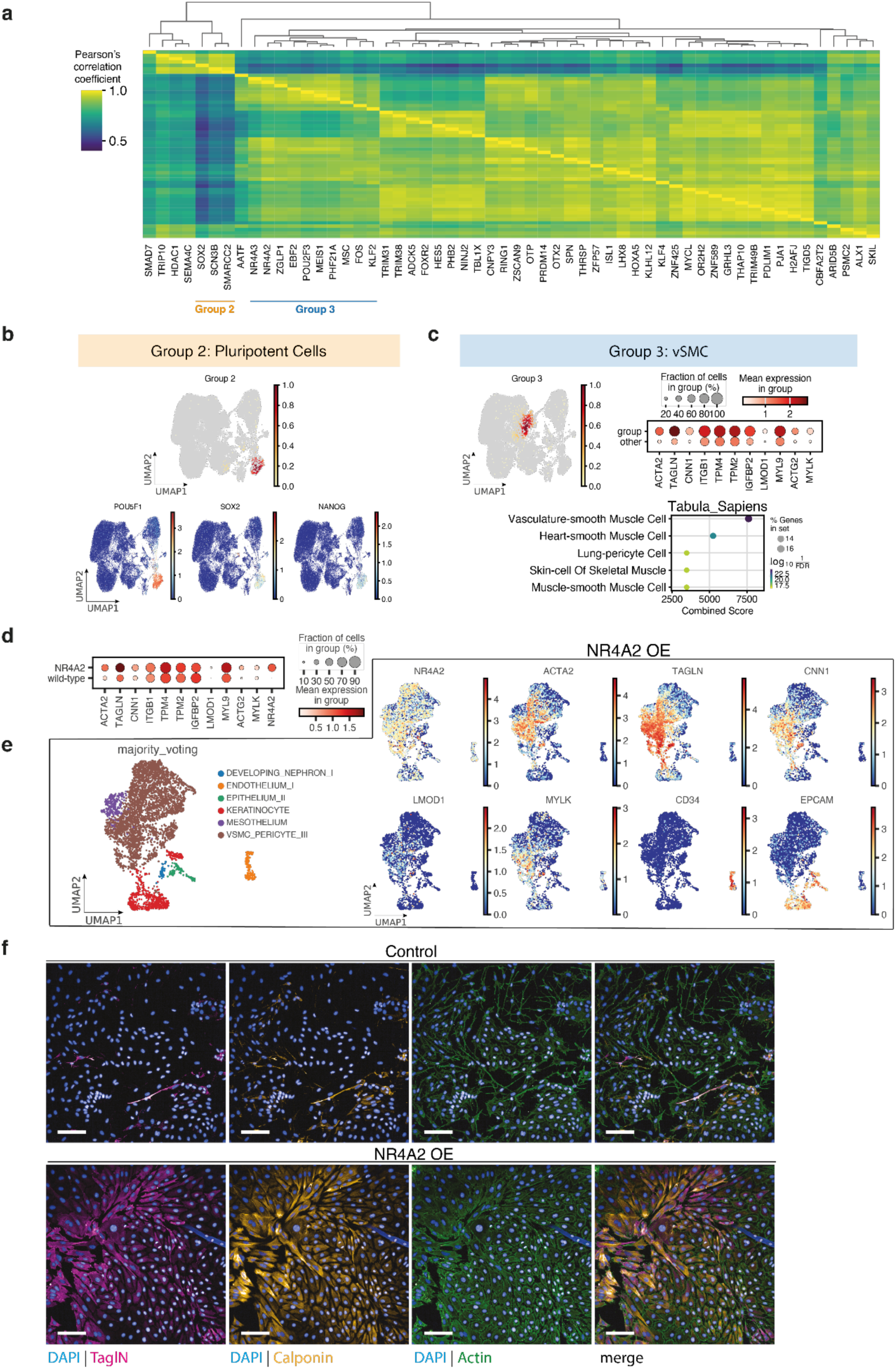
Groups of factors causing similar effects and validation of *NR4A2*. **a,** Heatmap showing pairwise Pearson correlations between TF-level average gene expression profiles from the D18 pooled screen. Columns and rows represent TFs, ordered by hierarchical clustering. The colour scale indicates Pearson correlation coefficients. **b,** Characterisation of Group 2 factors identified in **a**. Density plot showing the distribution of D18 cells assigned to Group 2 factors on the D18 pooled-screen UMAP embedding (same embedding as in Fig. 5d) (top). Bottom UMAPs show expression of pluripotency-associated marker genes in the same embedding. **c,** Characterisation of Group 3 factors identified in **a**. Density plot showing the distribution of D18 cells assigned to Group 3 factors on the D18 pooled-screen UMAP embedding (same embedding as in Fig. 5d) (top left). Dot plot showing expression of vSMC-associated marker genes in cells assigned to Group 3 factors (group) compared with all other D18 cells (other) (top right). Dot size indicates the percentage of cells expressing each gene, and dot colour indicates mean gene expression within each cell group. The bottom panel shows Enrichr analysis of genes upregulated in Group 3-assigned cells compared with other D18 cells. Differential expression was tested using the Wilcoxon rank-sum test, and the top 50 upregulated genes with *P*-value < 0.05 were tested against Tabula Sapiens-associated gene sets. **d,** Dot plot showing expression of vSMC-associated marker genes and *NR4A2* in cells harvested at D18 from *NR4A2*-overexpression and wild-type cultures. Dot size indicates the percentage of cells expressing each gene, and dot colour indicates mean gene expression within each cell group. **e,** UMAP embedding of cells harvested at D18 from *NR4A2*-overexpression cultures, coloured by CellTypist majority-voting predictions using the Pan Fetal Human model^43^ (left) and expression of selected vSMC, endothelial and epithelial marker genes (right). **f,** Immunostaining image of wild-type control and *NR4A2*-overexpressing cells harvested at Day 18 stained for DAPI and vSMC marker proteins (scale bar: 100 µm).

A second cluster of perturbations (Group 3) was characterised by increased expression of genes associated with vascular smooth muscle cells (vSMCs) (Fig. 6a and c). Cells assigned to factors within this group exhibited elevated expression of canonical smooth muscle markers including *ACTA2*, *TAGLN*, *CNN1* and *MYLK* (Fig. 6c). Consistent with these observations, genes upregulated in Group 3 cells relative to the remainder of the Day 18 population were strongly enriched for vasculature smooth muscle cell signatures from the Tabula Sapiens reference atlas^50,51^ (Fig. 6c). Together, these results demonstrate that perturbation-level correlation analysis can identify groups of genes that converge on related cell states and biological programmes.

Among the factors within the vSMC-associated group, NR4A2 induced one of the strongest smooth muscle transcriptional signatures (Extended Data Fig. 8e), and was therefore selected for further validation. *NR4A2* is best characterised for its context-dependent functions in neuronal development, particularly in post-mitotic midbrain dopaminergic neurons, but is expressed across multiple cell types and tissues. Notably, analysis of endogenous expression in the adult human brain single-nucleus atlas of Siletti et al. (2023)^52^ also identifies NR4A2 as cell-type enhanced in vascular-associated smooth muscle cells. Examination of cells from independently generated NR4A2-overexpression compared to wild-type revealed robust expression of smooth muscle markers including *ACTA2*, *TAGLN*, *CNN1* and *MYLK* (Fig. 6d and e). Automated cell type annotation using CellTypist^34^ further classified the majority of NR4A2-overexpressing cells as vascular smooth muscle cell/pericyte-like populations (Fig. 6e). To validate this phenotype, we repeated the differentiation using NR4A2 alone and assessed expression of smooth muscle markers by immunofluorescence. Consistent with the single-cell transcriptome results, NR4A2 overexpression induced a pronounced smooth muscle-like morphology accompanied by strong expression of TAGLN, calponin (encoded by *CNN1*) and filamentous actin compared with control cultures (Fig. 6f). Thus, rather than implying that NR4A2 is a lineage-specific determinant of vSMC identity, these results demonstrate that NR4A2 is sufficient to promote a vSMC-like transcriptional and morphological programme in the cellular context examined here. This is consistent with the broader context dependence of developmental TF activity, whereby transcriptional output is shaped by developmental state, chromatin accessibility and the regulatory environment in which a TF is expressed^53,54^. Together, these results demonstrate that pooled ORF screening can be used not only to identify individual lineage-inducing factors but also to systematically group perturbations according to their transcriptional outputs. This analysis identified groups of TFs converging on related cell states and revealed a previously unanticipated capacity of NR4A2 to promote a vSMC-like programme in differentiating hiPSCs.

## Discussion

Overexpression screening has lagged behind loss-of-function approaches due to technical challenges, despite the central role of gene expression in development, cellular reprogramming and cell engineering. This data will be critical for the generation of reliable virtual cell models and to drive cellular state changes for cell modelling and therapeutic development. STEER-seq addresses this challenge by combining site-specific genomic integration with pooled ORF libraries and barcode-enabled single-cell molecular readouts. By expressing all perturbations from a common genomic locus following Bxb1-mediated cassette exchange, STEER-seq minimises variability arising from integration site, copy number and promoter context while maintaining compatibility with pooled screening formats and avoiding the complexities of viral production. Although individual components of this strategy, including Bxb1-mediated library integration and defined landing pads, have important precedents, their combination here enables large-scale gain-of-function screening in hPSCs and during differentiation with direct single-cell transcriptomic and multimodal readouts. Consistent with this, direct comparison with the lentiviral TF screen of Joung et al.^4^ showed substantially stronger ORF overexpression with STEER-seq, which remained evident after 18 days of differentiation. The generation of both single and dual landing pad cell lines further extends this framework to controlled pairwise perturbation screening.

Application of STEER-seq to targeted and large-scale pooled screens identified regulators capable of directing diverse cell-state transitions under cytokine-free conditions. A focused screen of candidate haematopoietic regulators identified SPI1 as a potent inducer of haematopoietic differentiation, with expression of SPI1 alone promoting an HSC/MPP-like transcriptional state from hPSCs in the absence of exogenous haematopoietic cytokines. The inducible configuration further allowed us to separate induction of a cell state from its subsequent maintenance by withdrawing SPI1 after establishment of the progenitor-like population. Scaling the platform to large pooled ORF libraries further enabled identification and validation of regulators associated with distinct lineage programmes, including SMAD7-induced neural progenitor-like states, NR4A3-associated melanocyte differentiation and an NR4A2-induced vascular smooth muscle-like programme. These examples illustrate the ability of pooled gain-of-function screening to uncover factors sufficient to access diverse cellular states. By avoiding multi-step lentiviral production, the STEER-seq platform streamlines these experiments as well as downstream arrayed validation, and avoids biases associated with viral packaging. Targeted integration provides a common promoter, copy number and genomic context across perturbations, reducing variability introduced by position effects. Although transgene expression decreased during prolonged differentiation, robust overexpression remained detectable at D18. More broadly, these findings demonstrate how systematic perturbation screening can complement increasingly comprehensive single-cell reference atlases by providing direct functional validation of candidate regulators of cell state transitions.

An important consideration is that STEER-seq is designed to identify which fate a TF is sufficient to drive from a defined cellular starting state, rather than to reproduce endogenous developmental trajectories. This distinction is particularly relevant in hPSCs, where robust TF expression may be required to overcome the pluripotency programme and initiate differentiation. Previous comparisons of CRISPRa and ORF overexpression in hPSCs similarly showed that ORF expression of NEUROD1 or NEUROG2 induced neuronal differentiation under conditions where endogenous CRISPRa-mediated activation was insufficient. The phenotype induced by a TF will also depend on the chromatin landscape, cofactors and developmental competence of the recipient cell. Thus, context-dependent effects such as the NR4A2-induced vascular smooth muscle-like programme should be interpreted as demonstrating TF sufficiency in the cellular context examined, rather than implying an equivalent endogenous role during physiological development. Establishing the physiological relevance and endogenous mechanisms of individual hits will require subsequent mechanistic studies.

A distinguishing feature of STEER-seq is its compatibility with multimodal profiling technologies. The 3′ barcode architecture introduced here enables recovery of perturbation identity in joint RNA–chromatin accessibility assays, allowing transcriptional responses to be linked directly to chromatin remodelling within the same cells. This capability facilitated identification of candidate chromatin-opening factors and enabled reconstruction of enhancer-driven gene regulatory networks downstream of transcription factor induction. More generally, STEER-seq establishes a screening-to-mechanism workflow in which candidate regulators identified through pooled screening can be linked directly to downstream regulatory circuitry and chromatin-state changes. Together with the dual landing-pad architecture, this provides a framework for investigating both individual regulators and combinations of factors that cooperate or antagonise one another to establish cell identity.

As with any gain-of-function approach, STEER-seq has limitations that define how its results should be interpreted. Strong CAG-driven expression does not reproduce endogenous TF dosage or temporal regulation, but is a deliberate feature for testing TF sufficiency and engineering cell states. Transgene expression also decreases during prolonged differentiation, emphasising that no genomic locus can be assumed to remain equally active across all cellular states. Because different genomic contexts may perform differently across differentiation trajectories, we have generated additional landing-pad lines at new safe-harbour loci on chromosomes 4 and 7 for future evaluation. The current homotypic att-site configuration can also be further improved through heterotypic att sites, enhanced Bxb1 variants and optimised nuclear-localisation sequences to increase cassette-exchange efficiency and specificity, reducing total cell numbers needed for larger screens.

Several opportunities exist to further increase the scale and versatility of the platform. Although STEER-seq supported large-scale pooled ORF screening in the present study, future applications to for instance combinatorial screens will likely be constrained primarily by sequencing depth and the cost of high-resolution single-cell profiling. The 3′ barcode architecture broadens compatibility with a wide range of transcriptomic technologies and will facilitate implementation in high-throughput combinatorial indexing approaches, enabling substantially larger perturbation libraries and cell numbers to be screened simultaneously. Integration with more high-throughput single cell technologies such as 10x Genomics FLEX or Parse will further improve sensitivity for rare cell populations identified in large-scale screens, and enable more combinations of factors to be screened. Because perturbation identity is encoded within transcribed barcode sequences, future adaptations may also be compatible with emerging spatial transcriptomic technologies capable of capturing exogenous transcripts. Continued development of recombinase technologies should further expand integration efficiency, targeting flexibility and cargo capacity, providing opportunities to extend the platform beyond TF libraries to other coding perturbations and increasingly complex perturbation designs. By complementing rather than replacing existing lentiviral- and CRISPR-based screening approaches, STEER-seq provides an alternative when controlled copy number, genomic context and robust transgene expression are particularly important. Together, STEER-seq establishes a quantitative framework for pooled gain-of-function screening that integrates precise genomic engineering with transcriptomic and chromatin-level readouts. To support adoption by other groups, we provide the plasmids required to generate STEER-seq landing pad lines and donor vectors, together with a tutorial for the end-to-end computational analysis pipeline. By connecting controlled gain-of-function perturbation with single-cell transcriptomic and regulatory phenotyping, STEER-seq provides a route from discovery of factors capable of altering cell identity to mechanistic interrogation and rational engineering of the states they induce.

## Data Availability

The plasmids required to generate STEER-seq landing pad lines and donor vectors will be made available on Addgene upon acceptance of the manuscript. Plasmid maps and sequences are available in supplementary information.

The annotated integrated sc-RNA-seq and multiome objects for this study will be made available upon acceptance of the manuscript and can be explored online at https://cellxgene.cziscience.com. This data will be available upon publication on Zenodo at https://doi.org/10.5281/zenodo.21510149 along with raw images, flow cytometry, long read sequencing, digital PCR data and code. All newly generated sequencing data for this study were uploaded to ENA under accession codes EGAS00001005249 and EGAS00001007821, with sample accession numbers and details listed in Supplementary Table 1. All code scripts and notebooks for analysis presented in the manuscript are available at https://github.com/Teichlab/STEER-seq, with tutorials available at https://github.com/Teichlab/STEER-seq/blob/main/tutorials/github_tutorial.md. All code scripts and notebooks used for the analyses presented in this manuscript have been deposited, together with tutorials, in the reviewer-accessible Zenodo record above. As the findings described in this study are the subject of a pending patent application, the GitHub repository (https://github.com/Teichlab/STEER-seq) will be made publicly available upon acceptance of the manuscript. Private access to the GitHub repository can be provided to editors and reviewers upon request.

This publication is part of the Human Cell Atlas (www.humancellatlas.org/publications).

## Acknowledgements

We would like to thank the flow cytometry, sequencing and core informatics groups at the Sanger Institute for their support and members of the Bassett, Teichmann and Garnett labs for helpful discussions.

## Materials and methods

### hiPSC culture

Kolf_2_C1 is a male hiPSC line derived by the Human Induced Pluripotent Stem Cell Initiative (HipSci) consortium (www.hipsci.org) and subsequently single cell cloned (Kolf_2_C1, https://hpscreg.eu/cell-line/WTSIi018-B-1). Kolf_2.1S is an edited derivative of Kolf_2_C1, corrected for a 19 bp deletion in one copy of ARID2 by CRISPR, which has undergone extensive characterization. hiPSC lines were cultured under feeder-free condition in Stemflex medium (combo kit, Gibco TM A3349401) on Vitronectin substrate (VTN). VTN was used at 1:100 dilution of a 1 mg/ml stock solution in PBS, using 1 ml per six-well plate for coating. Incubate at RT for 1 h before aspirating and replacing with culture media immediately. After initial thaw, cells were clump-passaged 1:10 every 4–5 days and cultured in a humidified incubator at 37°C and 5% CO_2_.

### Generation of single and dual landing pad lines

For each nucleofection, 2 µg of landing-pad plasmid (see supplementary) per 1 × 10⁶ cells was added to pre-assembled Cas9–sgRNA ribonucleoprotein (RNP) complexes immediately prior to electroporation using a 4D-Nucleofector (Lonza), program CA-137. Guide RNAs were resuspended at 200 µM in IDT duplex buffer and used at a final concentration of 45 µM. RNPs were assembled by incubating 5 µL sgRNA (225 pmol) with 5 µL Cas9 protein (20 µg; ∼120 pmol) at room temperature for 10–20 min.

Actively growing hiPSCs (3–5 days post-passaging, ∼70% confluence) were washed with PBS, dissociated with Accutase for 6 min at 37 °C, and resuspended in StemFlex ™ supplemented with CloneR2. Cells were triturated to a single-cell suspension, counted, and 1 × 10⁶ cells were pelleted and resuspended in 100 µL P3 buffer (Lonza). Landing-pad plasmid was added to the RNP complexes immediately prior to mixing with the cells. Cells were transferred to nucleofection cuvettes and electroporated. Electroporated cells were gently recovered in StemFlex ™ + CloneR and plated on VTN-coated plates. Medium was exchanged after 24 h for StemFlex ™ without CloneR, and cells were fed daily until ∼70% confluence. Blasticidin selection (100 µg mL⁻¹) was initiated 48–72 h post-nucleofection. mApple fluorescence was monitored, and after ∼7 days of selection, cells were clonally expanded via FACS, seeding both low- and high-expressing populations to isolate single- and dual-landing pad lines.

A monoallelic landing pad line integration line was identified through PCR screening - junction PCRs were targeted to chromosome-insert boundary and were combined with wild-type allele PCR. The wild-type allele PCR product was sequenced by Sanger sequencing. A SNP proximal to the PAM of the sgRNA used for integration was found. A new guide was ordered and the monoallelic line was re-targeted in order to generate a bi-allelic landing pad integration line. The same PCR screening was performed – loss of wild-type allele PCR product was used for selecting bi-allelic integration candidates. Digital PCR was then used to verify the copy number of the landing pad in the lines (see below - Droplet digital PCR (ddPCR)).

Long range PCR amplification using wild-type allele primers was performed using Takara Ex Premier DNA polymerase and adjusted extension times to allow for amplification of the allele(s) containing the landing pad integration. The resulting PCR products were purified and sequenced using ONT - using random fragmentation with Rapid Sequencing Kit by Genewiz Azenta and using Ligation Sequencing Kit by Plasmidsaurus. The plots of coverage of the alleles of the monoallelic line are shown in Extended Figure 1.

### Droplet digital PCR (ddPCR)

Five to twenty nanograms of genomic DNA was used per 20 μl reaction including 10 μl of ddPCR Supermix for probes (no dUTP) (Bio-Rad, cat. #186-3023) and 0.5 μl of VIC-tagged TaqMan copy number reference assay, human RNase P (Thermo Fisher Scientific, cat. #4403326) as a reference. 2 pmol each of forward and reverse primers as well as 4 pmol of a probe for the target locus were used per 20 μl reaction. All the primers and probes used for droplet digital PCR (ddPCR) are listed in Supplementary Table 2. Droplets were generated using a QX200 Droplet Generator (Bio-Rad, cat. #1854002) and transferred to 96-well plates. PCR was performed with the following steps: 95°C for 3 min, 40× (95°C for 20 s and 60°C for 1 min), and 98°C for 10 min. The resulting fluorescein amidite (FAM) and 2’-chloro-7’-phenyl-1,4-dichloro-6-carboxy-fluorescein (VIC) fluorescence was read using a QX200 Droplet Reader (Bio-Rad, cat. #1864003) and positive droplets were determined. The copy number of the target locus was calculated as the ratio of the positive droplets for the target locus to those of the reference gene, multiplied by 2 (copy number of the reference gene).

### Cloning of donor plasmids for targeted HSC screening

Transcription factor coding sequences were synthesised as double-stranded DNA fragments (GeneWiz), with a unique barcode positioned at the 5′ end of each construct. Synthesised fragments were cloned into an attB-containing donor plasmid by restriction enzyme digestion and ligation. Each donor cassette comprised a 5′ barcode, Kozak consensus sequence, transcription factor open reading frame, P2A peptide and puromycin-resistance gene, flanked by attB recombination sites. The donor plasmid backbone contained an ampicillin-resistance gene for bacterial selection. Following transformation into competent *Escherichia coli*, individual colonies were selected on ampicillin-containing agar plates, and plasmid DNA was isolated and sequence-verified before use in pooled screening experiments. A complete list of transcription factor sequences, barcode sequences and corresponding donor constructs is provided in Supplementary Table 2.

### Generation of the inducible SPI1 donor plasmid

To generate the inducible SPI1 donor construct, the SPI1 coding sequence was subcloned into an inducible attB donor plasmid by restriction enzyme digestion and ligation. The donor cassette comprised two expression units flanked by attB recombination sites. The first expression cassette consisted of rtTA3G-P2A-puromycin. The second cassette contained a TRE (Tet-On) promoter driving expression of 5′ barcoded SPI1, followed by the woodchuck hepatitis virus post-transcriptional regulatory element (WPRE). Following Bxb1-mediated integration into the CLYBL landing pad, constitutive expression of rtTA3G enabled doxycycline-dependent induction of SPI1 expression from the TRE promoter. Plasmid maps and complete donor construct sequences are provided in Supplementary Table 2.

### High throughput barcoding of plasmid libraries

Gateway destination plasmid v2 was digested with BbsI-HF (NEB) and the large fragment was gel extracted. Primers containing a 12bp stretch of degenerate bases (random barcode) were used to amplify a 5’ UTR insert. The insert was cloned in the digested backbone using Gibson assembly. The resulting 5’ end barcode library was transformed into chemically competent OneShot ccdBr cells. Colony counts were performed to ensure sufficient barcode diversity.

Single colonies were also picked and sequenced using the Sanger method, these were then used for arrayed TF sub-cloning.

To insert 3’ barcodes in an arrayed format, vectors containing 5’barcode-TF pairs were digested with HindIII-HF(NEB). Oligonucleotides containing a 12bp barcode were cloned directly using Gibson assembly via the ssDNA cloning method.

### Cloning of donor plasmid library

Barcoded Gateway destination vector v2 was incubated in a Gateway LR cloning reaction with pools of TFs from the ORFeome collection. The pools were based on construct size according to the plate map of the provider (DNASU), aiming to ensure equimolar pooling. Pools of sub-cloned and barcoded TFs were transformed into bacteria separately and aliquots of the transformants were plated out to estimate colony counts. Final pooling was done by calculating the expected abundance of unique TFs per pool and appropriate aliquots were combined. The mixture was isopropanol precipitated and purified with the TXS method^56^.

### Long read sequencing

The library was processed following the PureTarget method (PacBio) and sequenced using PacBio Revio. The DNA was first dephosphorylated to remove potential adapter ligation sites. Two crRNAs were designed to target inside of the *bla* selectable marker in the Gateway destination vector v2. The crRNAs were complexed with tracrRNA, separately. RNP complexes with Sp.Cas9 were formed with the crRNA:tracrRNA The cloned TF library was digested with crRNA-Cas9 complexes targeting the backbone region. The crRNAs were designed with PAM-in orientation - Sp.Cas9 would remain bound to the drop-out fragment of length 113bp. The fragment containing the rest of the backbone and the barcode-TF pair varies in length but is at least 3950bp. The library was cleaned up using 1.2x SMRTbell beads, this step removes small fragments. The library was then dA-tailed with Taq polymerase. A hairpin adapter with 3’ dT overhangs and 5’ phosphate group was designed (see Supplementary Table 2). The adapter was resuspended in TE (pH 8.0) and diluted to 17μM in 10x annealing buffer. The adapter was heated to 95°C for 5min and then transferred to room temperature for 30min. The adapter was then ligated to the library following the PacBio protocol for PureTarget libraries (103-329-400 REV02). The library was further processed in accordance with this protocol and sequenced on a Revio sequencer machine.

### Donor plasmid integration and enrichment

Human induced pluripotent stem cell (hiPSC) landing-pad lines were maintained on vitronectin-coated plates in StemFlex medium (Thermo Fisher Scientific) at 37 °C and 5% CO₂. For donor integration, 1.8 × 10⁶ cells were reverse-transfected onto vitronectin-coated 10-cm plates with 15 µg of attB donor plasmid or donor plasmid library using Lipofectamine Stem Transfection Reagent (Thermo Fisher Scientific) according to the manufacturer’s instructions. Briefly, plasmid DNA was complexed with 80 µL Lipofectamine Stem in Opti-MEM, incubated for 10 min at room temperature, and added to the cells. StemFlex medium was added 2–4 h after transfection, and the medium was replaced 24 h later. To enrich for donor-integrated cells, puromycin was added at a final concentration of 0.6 µg mL⁻¹ beginning 24 h post-transfection and maintained for 24 h. Following selection, cells were either maintained for downstream analyses or transferred to APEL2 medium to initiate differentiation. For experiments using the doxycycline-inducible donor system, transgene expression was induced with 2 µg/mL doxycycline. For SPI1 differentiation experiments, doxycycline was added either from day 0 to day 18 or from day 8 to day 18, as indicated. Doxycycline was replenished at each medium change (every 3–4 days), and cultures were protected from light throughout the induction period. Doxycycline was withdrawn at day 18 for subsequent assessment of the induced cell state in the absence of continued SPI1 expression.

### Directed hematopoietic differentiation of engineered hiPSCs

Engineered hiPSCs carrying integrated transcription factor perturbations were differentiated using an embryoid body-based hemogenic endothelial differentiation protocol adapted from Ng et al. Briefly, cells were aggregated in ultra-low attachment 96-well plates in APEL medium, with or without cytokines, at 4,000 cells per embryoid body. For cytokine-supplemented conditions, day 0 medium contained BMP4 (20 ng ml−1), VEGF (25 ng ml−1), FGF2 (10 ng ml−1), SCF (25 ng ml−1), Activin A (10 ng ml−1) and ROCK inhibitor. On day 2, medium was partially refreshed with the same cytokines together with SB-431542 (3–4 μM) and CHIR99021 (3 μM). On day 4, medium was changed to APEL containing BMP4 (20 ng ml−1), VEGF (50 ng ml−1), FGF2 (10 ng ml−1), SCF (50 ng ml−1) and IGF2 (30 ng ml−1). On day 8, embryoid bodies were transferred onto growth factor-reduced Matrigel-coated plates in APEL containing VEGF (50 ng ml−1), FGF2 (10 ng ml−1), SCF (50 ng ml−1), IGF2 (20 ng ml−1), FLT3L (25 ng ml−1), TPO (25 ng ml−1) and IL-6 (25 ng ml−1), or maintained in matched cytokine-free medium where indicated. Cultures were refreshed on day 13 and harvested on day 18 for flow cytometry and single-cell profiling. The protocol was adapted from Ng et al. (2016). For arrayed validation experiments, individual transcription factors identified in pooled screens were integrated separately and differentiated under cytokine-free conditions using the same protocol. Cells were harvested at day 18, labelled using the 10x Genomics Fixed RNA Profiling (Flex) workflow and subjected to single-cell transcriptomic analysis, or analysed by immunofluorescence staining.

### Transcription factor overexpression and single-cell multiome profiling

hiPSCs carrying a single or dual landing pad architecture were engineered by site-specific Bxb1-mediated integration of transcription factor expression constructs, enabling controlled transgene insertion at defined genomic loci. Following puromycin selection, cells were cultured for 6 days in APEL medium in the absence of exogenous cytokines to allow establishment of transcription factor-driven cell state transitions. Cells were then harvested and processed for simultaneous single-cell chromatin accessibility and transcriptome profiling using the Chromium Single Cell Multiome ATAC + Gene Expression platform (10x Genomics) according to the manufacturer’s instructions. Perturbation identities were recovered from integrated transgene sequences, enabling joint analysis of transcription factor-induced changes in gene expression and chromatin accessibility.

### Immunostaining

For Immunostaining, differentiated cells were grown on 96-well PhenoPlates (6055300, Revvity) and fixed in 3.7% formaldehyde for 10 min at room temperature. Formaldehyde was stepwise replaced with PBS. Cells were permeabilised and blocked for 20 minutes in blocking solution (3% BSA, 0.3% Triton X-100 and 0.2 M Glycine in PBS). Meanwhile, 0.5 μg of primary antibodies (α-NES, ab242400 Abcam, RRID:AB_3678442; α-Calponin 1, ab46794 Abcam, RRID:AB_3073544; α-alpha smooth muscle actin, ab124964 Abcam, RRID:AB_11129103; α-TAGLN, ab14106 Abcam, RRID:AB_443021) were pre-conjugated with fluorophores using the Flexable 2.0 Antibody Labeling Kits (#KFA501, #KFA 502, #KFA503, proteintech). The conjugated antibody solution was diluted 1:10 in staining solution (1.5% BSA, 0.15% Triton X-100 and 0.1 M Glycine in PBS) to a total volume of 1ml and supplemented with 1 μg of DAPI (#62248, Thermo Fisher Scientific). 100 μl of staining solution was used per well and incubated at 4 °C overnight. Cells were washed three times in wash buffer (0.3 % Triton X-100 in PBS) and imaged in PBS using the Phenix Opera Plus HCS system.

### Flow cytometry analysis

Differentiated hiPSC cultures were harvested by collecting the culture supernatant to retain non-adherent cells, followed by dissociation of attached embryoid bodies with 1× TrypLE Select for 15–30 min at 37°C. Cells were diluted in calcium- and magnesium-free PBS, pelleted, resuspended in FACS buffer, and passed 1–3 times through a 23G needle to generate a single-cell suspension. Suspended and dissociated adherent fractions were recombined where appropriate, washed, and resuspended in FACS buffer. For HSC marker analysis, cells were stained with TrueStain Fc block, followed by incubation with fluorophore-conjugated antibodies against CD34, lineage markers, CD19, CD38, CD45RA and CD90 (Supplementary Table 2) for 30 min at 4°C. DAPI was added before acquisition to exclude dead cells. Samples were washed in FACS buffer (PBS 2% FBS) and resuspended prior to analysis. Flow cytometry data were acquired on XDP or Bigfoot flow cytometers and analysed using FCS Express.

### Single-cell RNA-sequencing experiment

For scRNAseq experiments, Chromium single cell 5’ reagent kits from 10x Genomics were used (Supplementary Table 1). Unsorted, or FACS-isolated cells were loaded onto each channel of the Chromium chip following the manufacturer’s instructions before droplet encapsulation on the Chromium controller. Single-cell cDNA synthesis, amplification, gene expression (GEX) and cell surface protein (CITE-seq) libraries were generated. Sequencing was performed on the Illumina Novaseq 6000 system. The gene expression libraries were sequenced at a target depth of 50,000 reads per cell. CITE-seq libraries were sequenced at a target depth of 5000 reads per cell.

### Barcode capture from 10x 5’ samples

To enrich for barcode capture of the barcode-TF pairs from the RNA-seq samples, a custom reverse transcription (RT) primer binding to the region of the 5’ UTR (downstream of the 5’ barcode) was spiked in immediately before GEM formation (Supplementary Table 2). Following normal GEM formation and reverse transcription, the cDNA was isolated from the GEMs. The low molecular weight fraction was also kept and purified as it contains most of the reverse transcribed barcode regions. A nested PCR was performed from this cDNA to amplify the barcode region. These barcode libraries were processed in parallel and sequenced using the same system as the rest of the RNA-seq library. In some experiments, only a nested PCR and no RT spike-in was performed and in others a specific RT primer was spiked into the 5’ 10x reactions.

### Targeted HSC screening and *SPI1* validation scRNA-seq analysis

#### Data preprocessing

Gene expression data were aligned using Cell Ranger (v7.2.0) against a modified GRCh38 reference (Ensembl release 2020-A) containing exogenous transgene sequences. To enable unambiguous detection of exogenous transcripts, the first 115 nucleotides of each transgene were appended to the reference FASTA. This region comprised an 8-nucleotide barcode unique to each exogenous TF, an 18-nucleotide Kozak/BamHI sequence, and the first 89 nucleotides of the TF coding sequence. This design ensured that any 90-nucleotide sequencing read mapping to the exogenous construct would contain at least one nucleotide unique to the transgene, thereby distinguishing exogenous from endogenous TF transcripts.

Low-quality cells were excluded based on the following criteria: fewer than 1,000 total UMI counts, fewer than 500 detected genes, mitochondrial transcript fraction greater than 20%, or a Scrublet^57^ (v0.2.3) doublet score greater than 0.4 or predicted doublet status.

For sample demultiplexing, each cell was assigned to the CITE-seq hashtag with the highest UMI count, provided that the dominant hashtag contained at least 15 UMIs and accounted for more than 50% of total hashtag counts within that cell.

Cells containing exogenous TF-derived reads were assigned to the TF with the highest exogenous UMI count, requiring that the dominant TF contributed more than 50% of all exogenous TF counts detected in the cell.

For dual landing-pad screening, barcode expression in non-cell droplets was assessed to estimate the distribution of barcodes arising from ambient RNA. Non-cell droplets were defined as droplets with total UMI counts ≤ 200. Across the two samples, the 95th percentile of non-zero barcode counts in non-cell droplets were all 1 UMI for all transgene barcodes, and we therefore defined positive barcode assignment as >1 barcode UMIs. For barcode calling, each cell was assigned to the barcode with the highest UMI count, provided that the dominant barcode contained at least 2 UMIs and accounted for more than 50% of the total barcode counts detected in that cell.

#### Data integration and annotation

Downstream analysis was performed using Scanpy^58^ (v1.11.4) following a standard workflow. Highly variable genes were identified using scanpy.pp.highly_variable_genes(batch_key=“file”), followed by principal component analysis with scanpy.tl.pca(), neighbourhood graph construction with scanpy.pp.neighbors(), and UMAP embedding via scanpy.tl.umap(). Leiden clusters identified from the integrated UMAP embedding were manually annotated based on canonical marker gene panels (Extended Data Fig. 4d). Cell type annotation was further supported by automated label transfer using CellTypist^34^ (v1.6.3) (Extended Data Fig. 4c) with the Pan_Fetal_Human.pkl reference model and majority voting enabled (celltypist.annotate(model=’Pan_Fetal_Human.pkl’, majority_voting=True)).

#### Enrichment testing

Enrichment of individual TF within specific cell populations was assessed using Fisher’s exact test. For each TF and target population combination, a 2×2 contingency table was constructed using:

1. the number of cells in the target population (endothelial cells at day 8 or haematopoietic cells at day 18) assigned to the given exogenous TF,
2. the number of cells in the same target population assigned to the EGFP control,
3. the number of cells in the baseline population (defined as day 0 iPSCs) assigned to the given exogenous TF, and
4. the number of baseline population cells assigned to the EGFP control.

Odds ratios and *P*-values were calculated using Fisher’s exact test, and *P*-values were corrected for multiple testing using the Benjamini-Hochberg procedure.

### Multiome data analysis

#### Preprocessing and differential gene expression analysis

For Multiome gene expression preprocessing, raw Cell Ranger ARC gene expression matrices were first subset to the Gene Expression modality and processed with CellBender^59^ ‘remove-background’. CellBender was run per sample using the Cell Ranger filtered cell count to define priors: ‘expected-cells’ was set to 60% of Cell Ranger-called cells, and ‘total-droplets-included’ was set to the Cell Ranger cell count plus twice this number, capped at 15,000 additional barcodes. Models were run with false-positive rates of 0.01, 0.05 and 0.1, using GPU acceleration; where required, ‘--low-count-threshold 1’ was added to handle samples with too few high-count empty droplets. Cell calling for Multiome libraries was further evaluated using EmptyDropsMultiome^60^ on the raw Cell Ranger ARC feature-barcode matrix.

Transcription factor barcode count matrices were analysed separately to assign perturbation identities. For each sample, barcode count matrices were split by barcode position, and each cell was assigned to the perturbation with the highest barcode count.

After initial cell calling and CellBender correction, Multiome gene expression datasets were combined and analysed in Scanpy. Counts were normalised, log-transformed, highly variable genes were identified, and PCA, nearest-neighbour graph construction, Leiden clustering and UMAP embedding were performed. Doublet filtering used a combination of gene-expression doublet scores, ATAC doublet scores and AMULET^61^ calls. Cells were overclustered at high Leiden resolution, cluster-level doublet statistics were calculated, and clusters with high consensus doublet scores were removed together with high-confidence AMULET doublets.

The filtered gene-expression object was reclustered and annotated using marker gene expression and perturbation identity. Perturbed iPSC cells were then retained for downstream analysis, normalised and log-transformed, and reprocessed with highly variable gene selection, PCA, neighbour graph construction, Leiden clustering and UMAP. Differential gene expression between perturbations, including comparisons against the control perturbation, was tested using Scanpy’s Wilcoxon rank-sum test.

#### Peak calling and differential chromatin accessibility analysis

For the Multiome ATAC modality, Cell Ranger ARC fragment files were processed with ArchR^62^ using the hg38 genome. Arrow files were generated per sample with ‘createArrowFiles’, requiring TSS enrichment ≥5, at least 1,000 fragments and no more than 100,000 fragments per cell. TileMatrix and GeneScoreMatrix generation were enabled, promoter regions were defined as 2,000 bp upstream and 100 bp downstream of transcription start sites, and chrY was excluded. ATAC doublets were assessed with AMULET using Cell Ranger ARC fragment files, barcode calls from the preprocessing workflow, human autosomes and hg38 RepeatMasker regions. Filtered gene-expression annotations were transferred to the ArchR project by matching sample-barcode identifiers. Cells lacking matched RNA-derived annotations were removed before downstream ATAC analyses. Pseudo-bulk accessibility profiles were generated with ‘addGroupCoverages’, grouping cells by perturbation, with at least two pseudo-replicates per group, 50–500 cells per replicate, and a sampling ratio of 0.8. Reproducible peaks were called with MACS2^63^ through ArchR using ‘addReproduciblePeakSet’, grouping by perturbation. Peaks were called with q-value filtering at 0.01, excluding chrY and chrMT, extending summits by ±250 bp, allowing up to 5,000 peaks per cell and up to 200,000 peaks per group. A cell-by-peak accessibility matrix was then generated with ‘addPeakMatrix’ using a ceiling of 5 without binarisation. ATAC quality was assessed using ArchR fragment-size distributions and TSS enrichment profiles across samples.

Differential chromatin accessibility was tested from the ArchR PeakMatrix using ‘getMarkerFeatures’, grouping cells by perturbation and correcting for TSS enrichment and log10-transformed fragment counts. Marker peaks were identified using a t-test, and regions with FDR ≤0.2 and log2 fold change ≥0 were retained.

#### GRN inference using SCENIC+

Gene regulatory networks (GRNs) were inferred from paired single-cell RNA-seq and ATAC-seq data using the SCENIC+^29^ framework. To reduce sparsity, cells were first aggregated into metacells using SEACells, which was run separately for each perturbation. The number of metacells was determined as approximately one metacell per 20 cells, using the UMAP embedding to construct the similarity kernel and 10 waypoint eigenvectors for archetype initialisation. Metacell quality was evaluated using perturbation purity, compactness and metacell size, and metacells containing fewer than 10 cells were excluded from downstream analyses. RNA and ATAC profiles were summarised for each metacell and combined into a MuData object^64^.

Chromatin accessibility topics were identified using pycisTopic^65^ after restricting analyses to peaks accessible in at least 5% of cells in at least one perturbation group. Topic models were binarised using both Otsu thresholding and top-ranked regions, and differential accessible regions identified by ArchR were incorporated as complementary regulatory region sets.

Transcription factor motif enrichment analysis was performed using pycistarget on cisTopic-derived topics and ArchR differential accessibility regions. Motif enrichment was evaluated against custom cisTarget ranking and score databases generated from the experiment-specific peak set, using motif annotations from the v10 non-redundant clustered motif collection. Promoter regions were excluded during motif enrichment analyses.

A SCENIC+ object was initialised by integrating the filtered gene expression AnnData object, the pycisTopic object and pycistarget motif enrichment results. GRNs were inferred using SCENIC+, linking transcription factors to target genes through accessible regulatory regions. Regulatory search windows extended from 1 kb to 150 kb upstream and downstream of each gene. To assess network robustness, SCENIC+ was repeated on 20 independent random subsamples, each containing 80% of cells. Standard SCENIC+ eRegulon filtering was applied to each run, and eRegulon metadata together with TF-to-gene and region-to-gene links were exported for downstream consensus analysis.

High-confidence eRegulons identified across independent SCENIC+ runs were combined into a consensus regulatory network and re-imported into the SCENIC+ object. Region-based and gene-based eRegulon activities were quantified using AUCell with an AUC threshold of 0.005. eRegulon activities were subsequently summarised across perturbations using pseudobulk profiles, and transcription factor–cistrome correlations were calculated to prioritise robust regulatory programmes. Downstream analyses included eRegulon activity visualisation, regulon specificity score (RSS) calculation and dimensionality reduction based on eRegulon activity profiles.

Differentially accessible regions identified by ArchR were used throughout the regulatory network inference and interpretation pipeline to support motif enrichment analyses and the identification of perturbation-associated regulatory programmes.

### Pooled screen scRNA-seq analysis

#### Data preprocessing and integration

Barcode-ORF libraries were sequenced using PacBio as described above. Sequencing reads were mapped with minimap2-2.28 using the map-hifi preset, using a set of CDS sequences generated from the 10x GRCh38 2020-A reference (GENCODE v32/Ensembl 98), to identify the TF associated with each barcode. Mapped reads were post-processed to identify the barcode based on its flanking sequences (CGCGCCGAAGCGACC and CTTGTCTCGCTCCGG), requiring a normalised indel similarity score of at least 90 to identify flanking sequencing hits. Barcodes were required to be between 10 and 14bp long, and barcode-ORF combinations with 10 or fewer reads were discarded. Barcodes linked to more than one ORF were annotated as ‘MULTIHIT’. The barcode mapping results are provided in Supplementary Table 3 and were used to generate the feature reference file (Supplementary Table 4) for CITE-seq analysis during Cell Ranger processing.

Gene expression libraries were aligned to the GRCh38 2020-A reference (GENCODE v32/Ensembl 98) using Cell Ranger v9.0.0. Barcode expression in non-cell droplets was assessed to estimate the distribution of barcodes arising from ambient RNA. Non-cell droplets were defined as droplets with total UMI counts ≤ 200, based on barcode rank plots from the Cell Ranger web_summary.html files for the four samples. These plots showed a second plateau, corresponding to the onset of empty droplets, at approximately 200 to 400 UMIs.

Low-quality cells and doublets were excluded using the same criteria described in the section “Targeted HSC screening and *SPI1* validation scRNA-seq analysis”. For barcode calling, each cell was assigned to the barcode with the highest UMI count, provided that the dominant barcode contained at least 6 UMIs and accounted for more than 50% of the total barcode counts detected in that cell.

Integration was performed using scvi-tools^46^ (v1.3.3). The top 2000 highly variable genes across all samples were selected using scanpy.pp.highly_variable_genes(n_top_genes=2000, flavor = “seurat_v3”, layer=”counts”, batch_key=“file”). We then trained an scVI model with n_layers = 2, n_latent = 30.

All CellTypist^34^ results shown were using CellTypist (v1.6.3) with the Pan_Fetal_Human.pkl reference model and majority voting enabled.

#### Comparison with the lentiviral-based platform

For comparison with a lentiviral-based TF overexpression platform, scRNA-seq data from Joung et al. 2024^4^ were downloaded from GEO (GSE216595). This dataset profiles hESCs transduced with a library of 90 TF ORFs and cultured for 7 days. To maximise comparability with the data generated in this study, only the 10x Genomics dataset was used, although it is worth to note that Joung et al. 2023 used the 10x Chromium Single Cell 3′ v2 platform, whereas the pooled screen in this study was profiled using the 10x Chromium Single Cell 5′ v3 platform.

Gene expression matrices were loaded from the mapped count matrices, and barcode-to-ORF assignments were obtained from GSM6681047_180124_TFmap.csv.gz. Counts were normalised to a total of 10,000 counts per cell and log1p-transformed. For each TF ORF, cells assigned to barcodes corresponding to that ORF were identified, and the mean log1p-normalised expression of the corresponding TF was calculated. Control cells were defined as cells with no barcode assigned. For each ORF, the degree of overexpression was quantified as the difference between the mean log1p-normalised expression in barcode-positive cells and the mean log1p-normalised expression in control cells. Only ORFs represented by at least 10 barcode-positive cells were included.

The same calculation was performed for the pooled RMCE screen generated using STEER-seq at D0 and D18. The resulting ORF overexpression distributions were compared with the Joung et al. lentiviral dataset in Fig. 5b.

#### Neighbourhood connectivity

All cells from the pooled screen harvested at D18 were selected, and the neighbourhood graph was recomputed using scVI latent embeddings. The resulting connectivity matrix (adata.obsp[‘connectivities’]) was used to quantify whether cells assigned to the same ORF were transcriptionally closer to one another than expected by chance.

For each ORF, cells assigned to that ORF by barcode calling were identified. ORFs represented by at least 20 cells were retained for analysis. For each retained ORF, the within-ORF mean pairwise connectivity was calculated from the neighbourhood graph by extracting the connectivity values between all pairs of cells assigned to that ORF and dividing the summed connectivity by n(n − 1), where n is the number of cells assigned to that ORF, thereby excluding self-connections.

To estimate the background level of connectivity expected by chance, the same number of cells was randomly sampled from all D18 pooled-screen cells without replacement, and the mean pairwise connectivity was calculated using the same approach.

#### Pearson correlation of average gene expression profiles

Data from the D18 pooled screen were used for this analysis. ORFs were included if they were assigned to at least 20 cells and had a mean within-ORF connectivity greater than 0.005, as calculated from the D18 neighbourhood graph described above. For each selected ORF, an average gene expression profile was generated by calculating the mean expression of each highly variable gene across all D18 cells assigned to that ORF. Pairwise Pearson correlations were then calculated between ORFs using these averaged highly variable gene expression profiles.

Hierarchical clustering was performed on the ORF-ORF Pearson correlation matrix to identify groups of ORFs with similar average gene expression profiles. Clusters were defined by cutting the dendrogram at a distance threshold of 0.5.

### Validation scRNA-seq analysis from 10x FLEX

Data were preprocessed and integrated as described in the section “Targeted HSC screening and *SPI1* validation scRNA-seq analysis”. Downstream analysis was performed using the standard Scanpy^58^ workflow.

CellTypist^34^ predictions were generated using CellTypist (v1.6.3) with either the Pan_Fetal_Human.pkl or Developing_Human_Brain.pkl reference model, as specified in the relevant figure legends and main text. Majority voting was enabled for all CellTypist analyses.

Additional cell type prediction with SCimilarity^47^ (v0.4.1) was performed following the tutorial (https://genentech.github.io/scimilarity/notebooks/cell_annotation_tutorial.html) using unconstrained annotation, allowing cells to be classified as any cell type represented in the SCimilarity reference.

