## Supplementary Figures for "STEER-seq: a recombinase based platform for pooled multimodal single-cell perturbation screening"

### Extended Data Figures

#### Extended Data Fig. 1

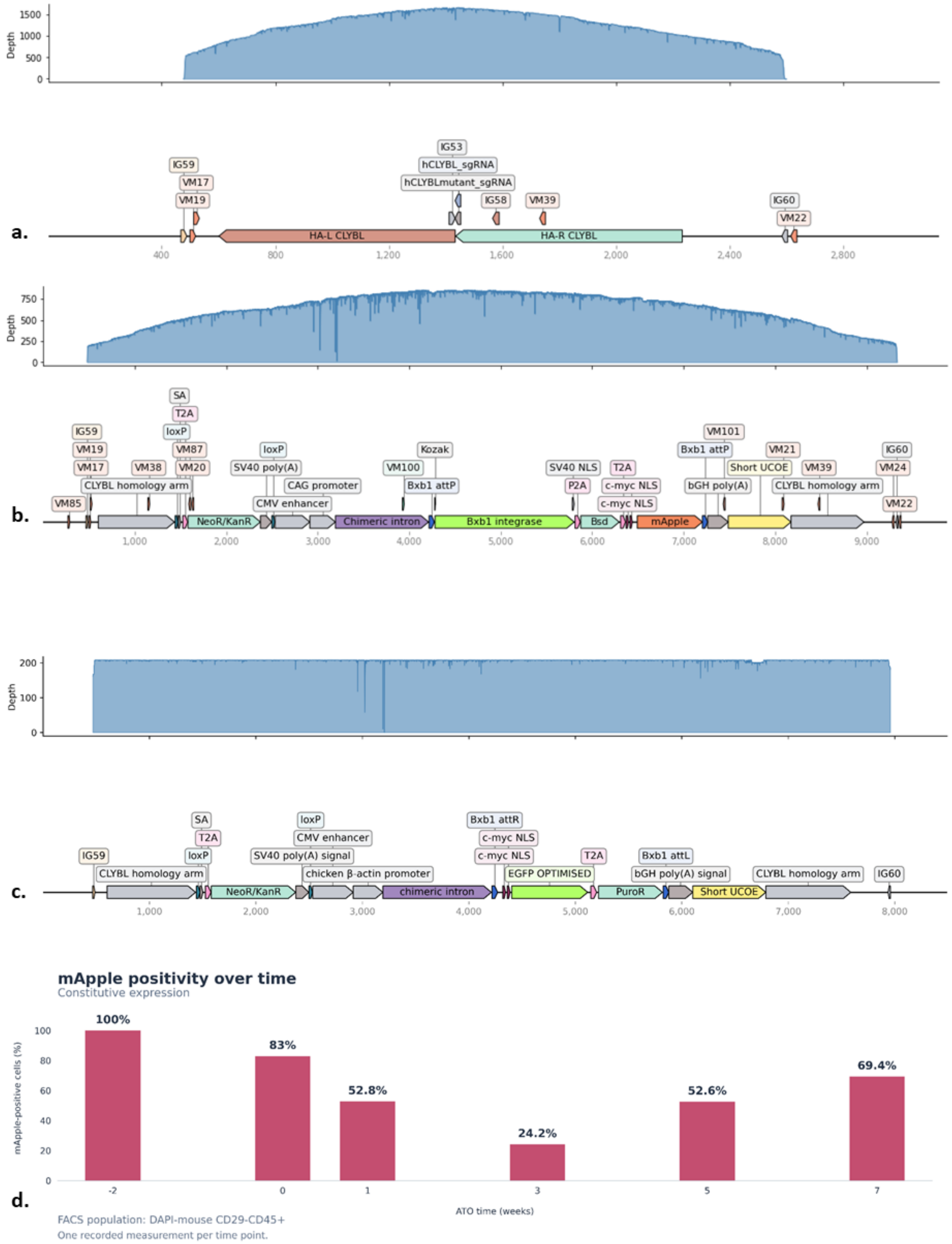

**Extended Data Fig. 1 | Long-read sequencing validation of landing pad integration at the CLYBL locus.**

**a,** Coverage plot from Oxford Nanopore long-read sequencing of the wild type allele following long-range PCR amplification using locus-specific primers. Purified amplicons were sequenced using the Oxford Nanopore Rapid Sequencing Kit (Genewiz Azenta), demonstrating the presence of an unmodified CLYBL allele **b,** Coverage plot of the corresponding engineered allele obtained from the same monoallelic landing pad cell line with the same locus-specific primers, confirming the expected landing pad structure and integration at the CLYBL locus. Amplification of both amplicons was done in a single pot (same primers) with Takara Ex Premier DNA polymerase using extended extension times to enable amplification across the landing pad insertion. Reads were mapped to a reference (fasta format) containing both alleles using minimap2.29-r1283[66] with the map-ont preset and then alignments were filtered for reads with MAPQ equal or above 20 before exporting coverage depth (both with samtools v1.23)[67,68] and plotting with DNA Features Viewer 3.1.3 and Matplotlib 3.3.4. The plotting script is in supplementary data. **c,** Coverage plot of the engineered allele post-transfection with RMCE donor containing EGFP (EGFP_Constitutive_pUC-GW-Amp) and selection with puromycin. The PCR product was sequenced using Oxford Nanopore Ligation Sequencing Kit (Plasmidsaurus). Reads were filtered on size (between 7kbp and 8kbp) and completeness (i.e presence of primer binding sites on both ends) and plotted against a reference map as above. **d,** Stability of CLYBL landing-pad expression during differentiation. Percentage of mApple-positive cells during haematopoietic differentiation of the unrecombined landing-pad hiPSC line in the artificial thymic organoid (ATO) system. mApple is constitutively expressed from the CAG promoter within the CLYBL landing pad and was monitored over 7 weeks of differentiation. Expression decreased during early differentiation but subsequently recovered and remained readily detectable at later time points.

**Extended Data Fig. 2**

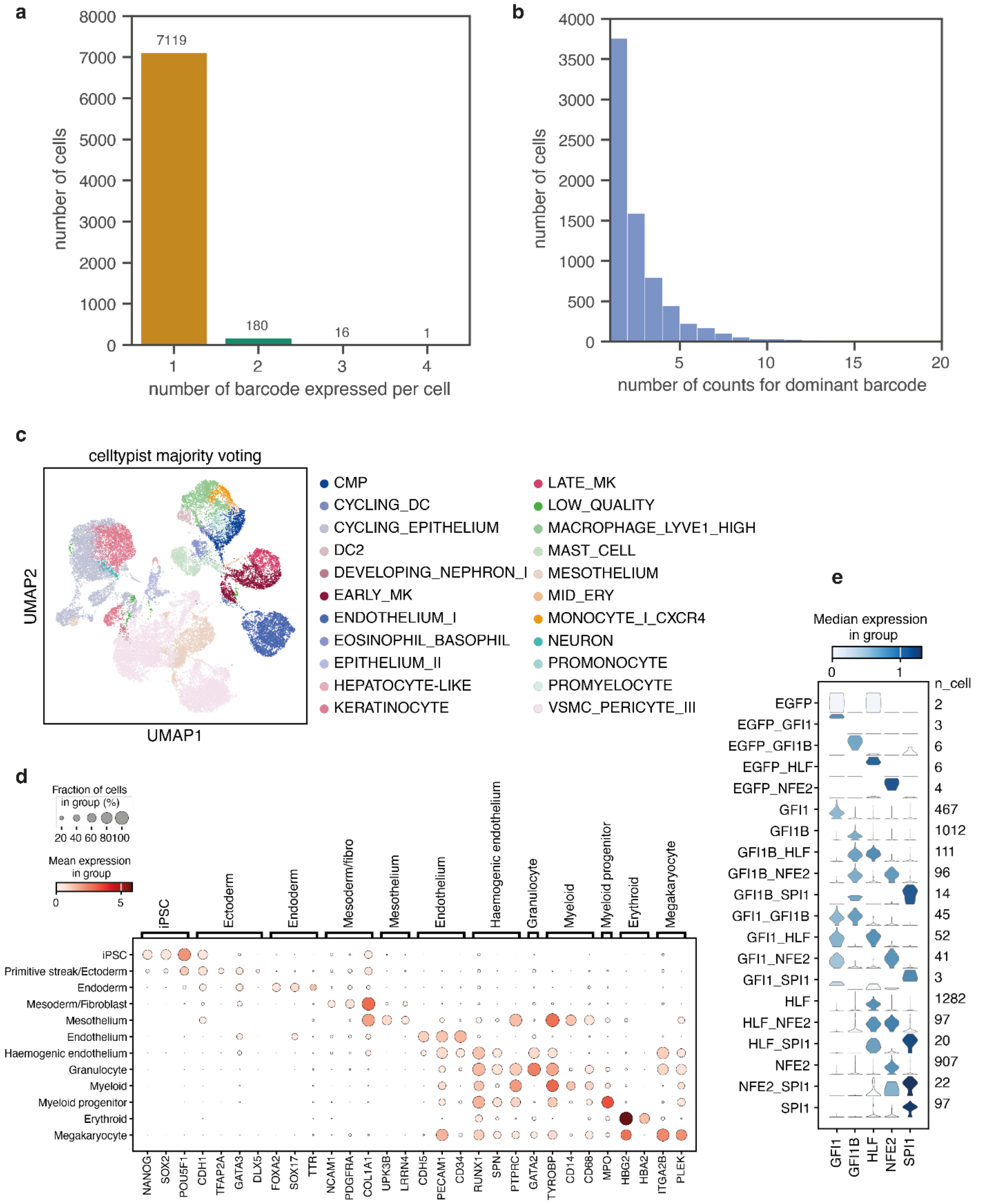

**Extended Data Fig. 2 | Targeted HSC screen**

**a,** Histogram showing the distribution of cells by the number of detected exogenous TF barcodes per cell. Numbers above bars indicate the total number of cells in each category. **b,** Histogram showing the distribution of cells by the UMI counts of the dominant exogenous TF barcode detected per cell. **c,** UMAP embedding coloured by predicted cell type annotations generated using CellTypist[34] with majority voting using the Pan Fetal Human model[43], shown on the same integrated embedding as in Fig. 2c. **d,** Dot plot showing the expression of selected marker genes across cell types identified in the integrated scRNA-seq dataset shown in Fig. 2c. Dot colour indicates mean expression level (log-normalised counts), and dot size represents the fraction of cells expressing each gene. Rows correspond to cell types and columns correspond to marker genes. **e,** Stacked violin plots showing log-normalised expression of each TF (*x*-axis) across cells assigned to the corresponding exogenous TF barcode combinations (*y*-axis). Cell numbers for each barcode combination are indicated on the right. Data is from dual landing-pad iPSCs transfected with a library containing five TFs and *EGFP*, harvested at D0 (2222 cells) and D18 (8231 cells).

#### Extended Data Fig. 3

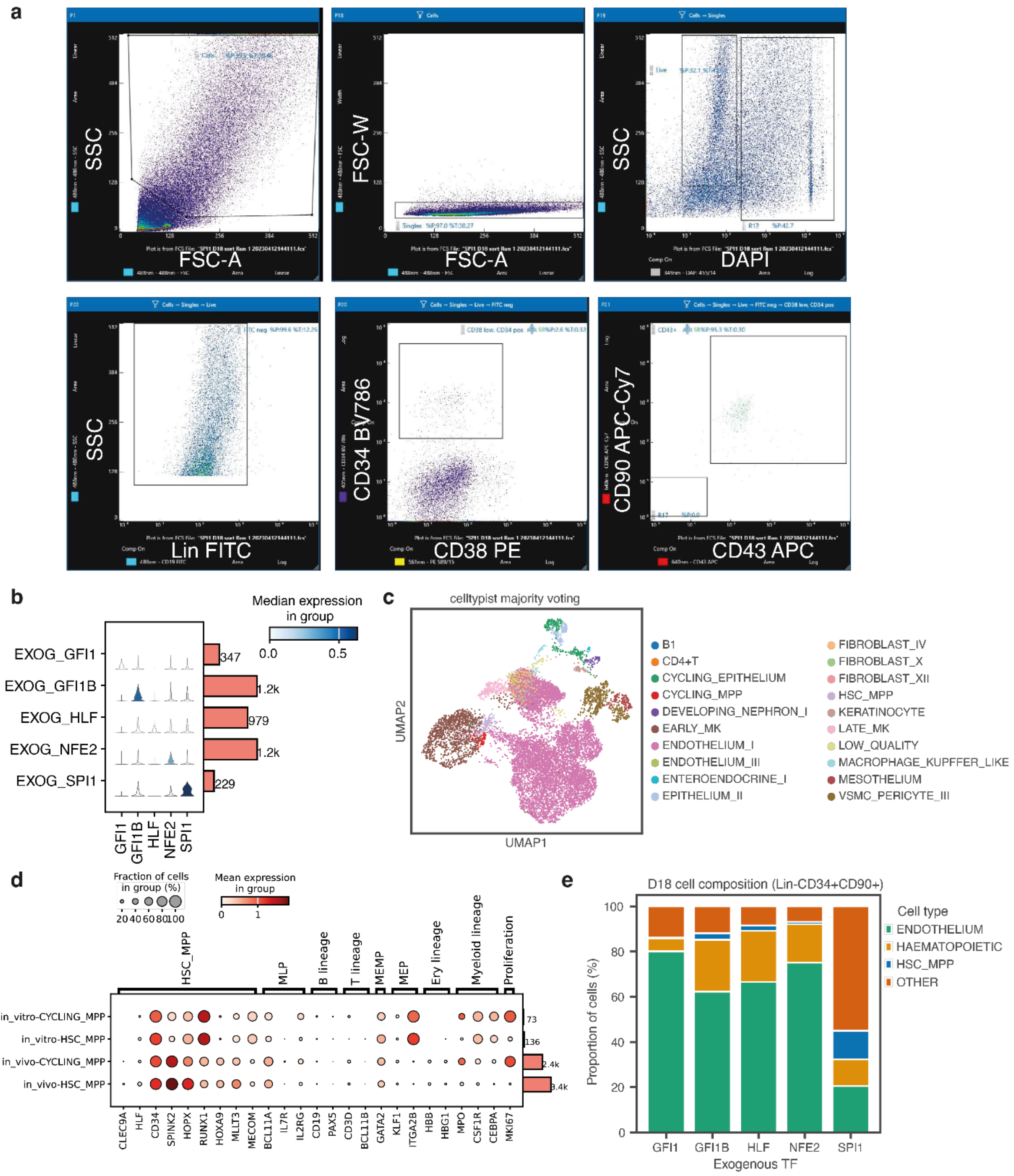

**Extended Data Fig. 3 | *SPI1* validation with constitutive promoter**
**a,** Representative flow cytometry plots showing the sorting strategy to enrich for HSPCs. Cells were sequentially gated for singlets, viability (DAPI−), lineage negativity (Lin−), and expression of CD34 and CD90. Lin−CD34+CD90+ cells were sorted and subjected to single-cell RNA sequencing. **b,** Stacked violin plots showing log-normalised expression of each TF across cells assigned to the corresponding exogenous TF barcode. **c,** UMAP embedding coloured by predicted cell type annotations generated using CellTypist[34] with majority voting using the Pan Fetal Human model[43]. **d,** Dot plot showing the expression of selected marker genes across *in vitro* and *in vivo* HSC/MPP populations, with *in vivo* reference data obtained from Suo et al. (2022)[43]. Dot colour indicates mean gene expression (log-normalised counts), and dot size represents the fraction of cells expressing each gene. Rows correspond to cell populations and columns correspond to marker genes. The total number of cells in each cell population is shown in the accompanying bar plot on the right. **e,** Stacked bar plots showing the predicted cell type composition of the D18 Lin−CD34+CD90+ sorted populations across different differentiation conditions from the scRNA-seq data in **c**.

#### Extended Data Fig. 4

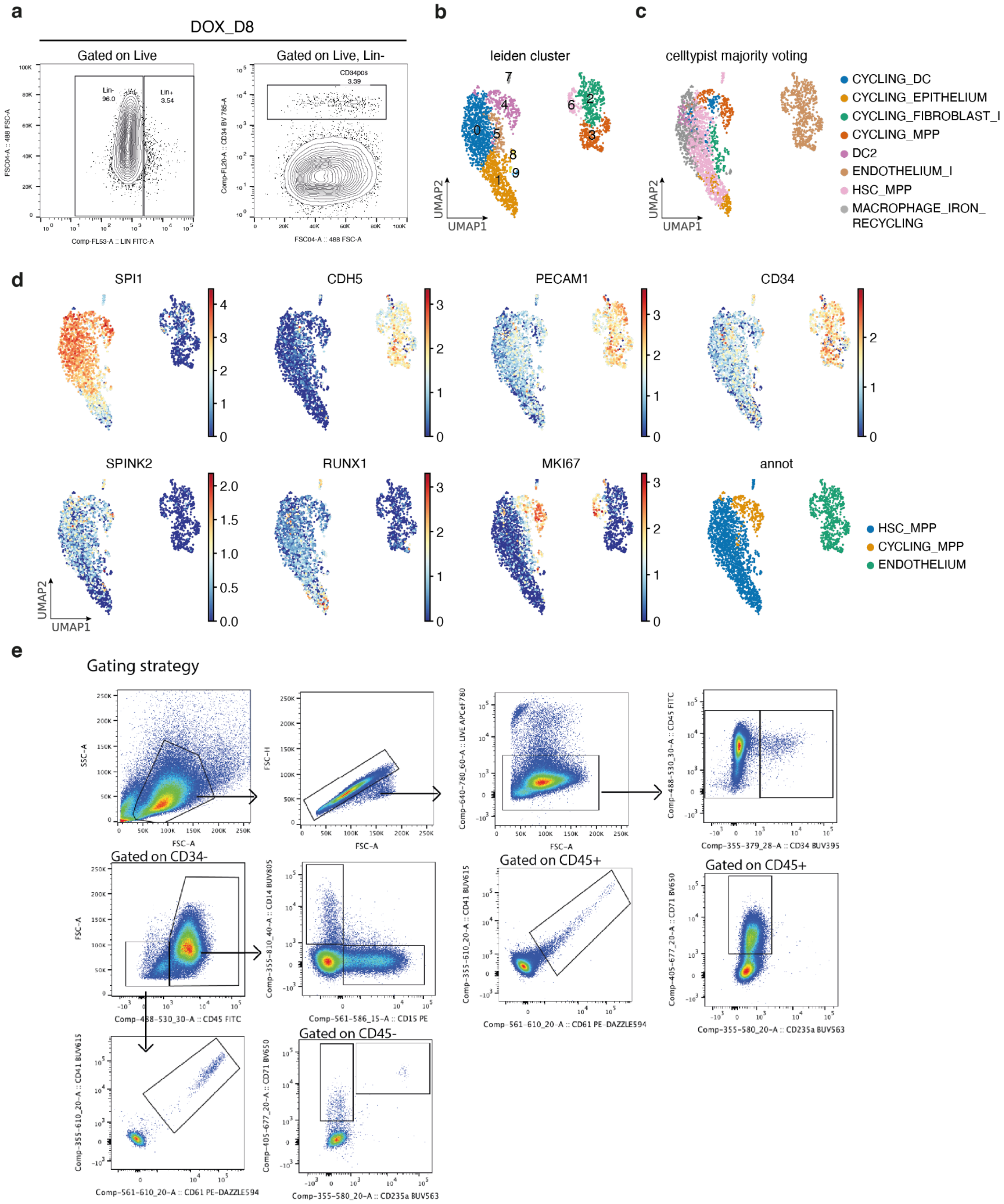

**Extended Data Fig. 4 | *SPI1* validation with inducible promoter**

**a,** Representative flow cytometry plots showing the sorting strategy used to enrich for HSPCs. Cells were harvested at D18 from inducible *SPI1* cultures treated with doxycycline between D8 and D18 (DOX_D8) and Lin–CD34+ cells were isolated for scRNA-seq analysis. **b–d,** UMAP embedding of scRNA-seq data from sorted D18 Lin–CD34+cells from inducible *SPI1* cultures (same embedding as Fig. 3d). Cells are coloured by Leiden cluster assignment (**b**), CellTypist majority-voting predictions using the Pan Fetal Human model[43] (**c**), and expression of selected marker genes together with the final manual cell type annotations (**d**). Gene expression values are shown as log-normalised counts. **e,** Representative flow cytometry plots showing the gating strategy used for the lineage differentiation assays presented in Fig. 3g. Cells were sequentially gated on live single CD34– cells, followed by lineage-specific marker combinations to identify megakaryocytic (CD45–CD41+CD61+), erythroid progenitor (CD45–CD71+), monocytic (CD45+CD14+), and granulocytic (CD45+CD15+) populations.

#### Extended Data Fig. 5

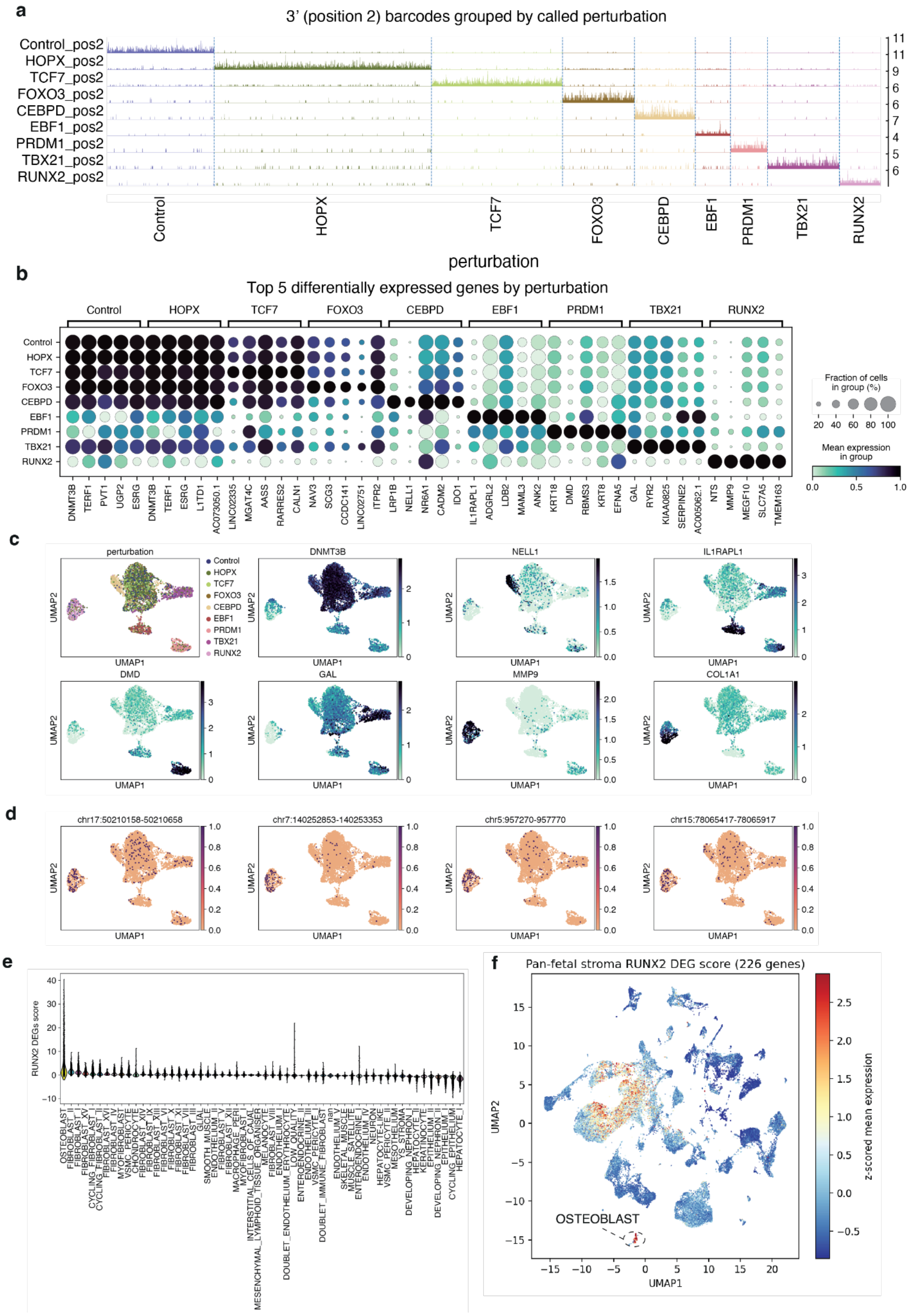

**Extended Data Fig. 5 | TF overexpression screening with single-nucleus multiome readout**
**a,** Track plot of TF barcode counts mapping to the 3' position (after TF cDNA sequence and ) in each nucleus grouped by the called perturbation based on the highest barcode count. Control indicates a construct containing the Venus fluorescent protein in the place of the TF. **b,** Dot plot of most differentially expressed genes between each perturbation an all other perturbations, with dot size indicating percent of nuclei expressing the gene and colour indicating the normalised expression. Differentially expressed genes are shown on the x axis for each overexpressed TF, which are indicated on the y axis. Genes on the x axis are ordered and grouped by perturbation. **c,** Gene expression-based UMAP representations with colour indicating perturbations and expression of selected differentially expressed genes. **d,** Gene expression-based UMAP representations with colour indicating accessibility of selected differentially accessible regions between RUNX2 overexpression and control. **e,** Expression of a score of differentially expressed genes (DEG) between RUNX2 overexpression and control across cell types in a foetal pan-tissue atlas. **f,** Expression of the RUNX2 DEG score visualised on a UMAP representation of the foetal pan-tissue atlas.

#### Extended Data Fig. 6

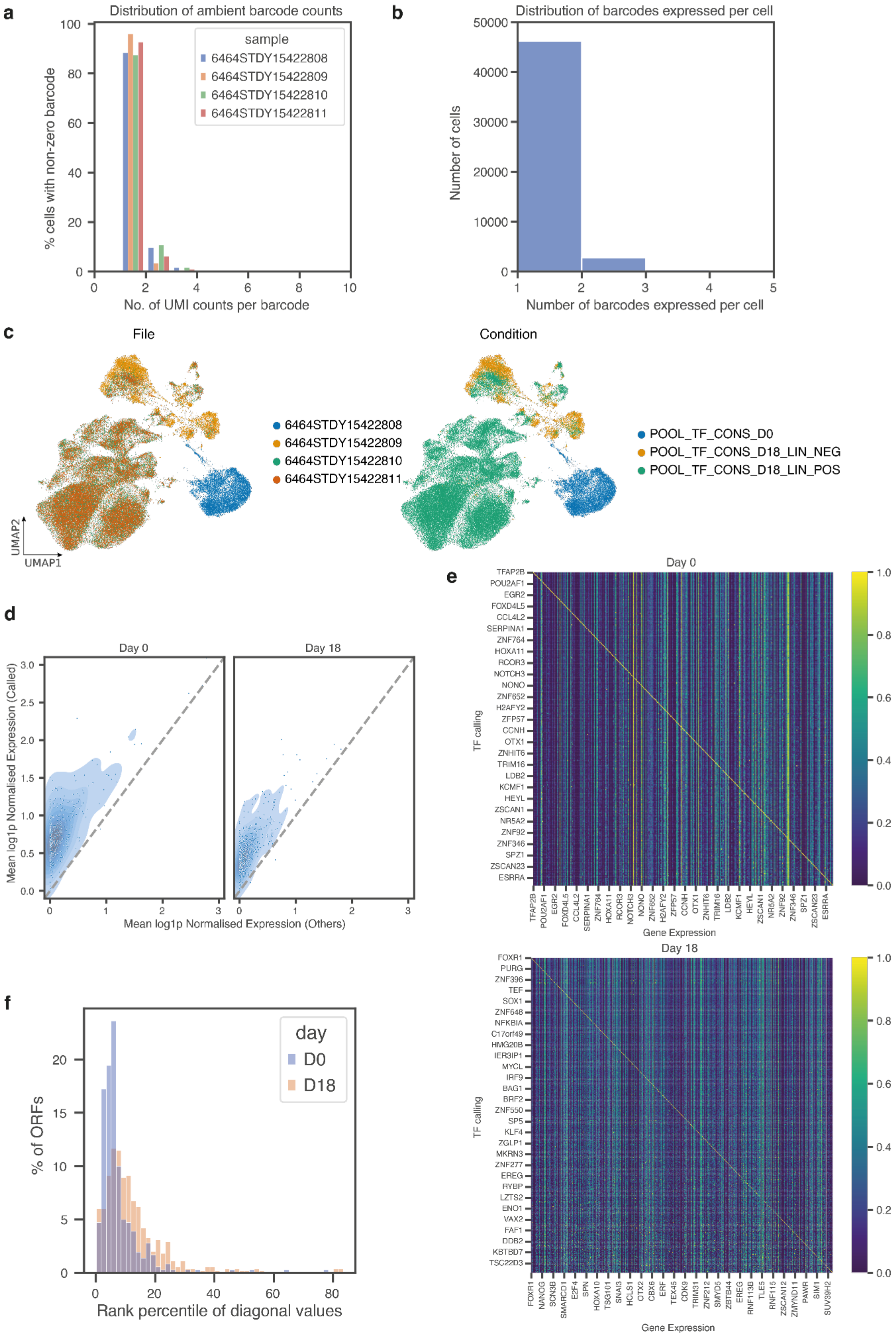

**Extended Data Fig. 6 | Barcode calling and validation of ORF overexpression in the pooled TF screen.**

**a,** Histogram showing the distribution of non-zero barcode UMI counts detected in non-cell droplets across the four samples. **b,** Histogram showing the number of unique barcodes detected per cell after applying the barcode UMI count threshold of > 5. **c,** UMAP embeddings of the integrated scRNA-seq dataset coloured by sample (left) or experimental condition (right). POOL_TF_CONS_D0 contains cells harvested at D0 of differentiation and sorted as mApple−, indicating successful recombination of the donor plasmid and loss of the original mApple signal from the landing pad. POOL_TF_CONS_D18_LIN_NEG contains D18 cells sorted as mApple−CD45−Lin−, whereas POOL_TF_CONS_D18_LIN_POS contains D18 cells sorted as mApple−CD45−Lin+ together with mApple−CD45+ cells. **d,** Mean expression of each ORF in cells assigned to any barcode linked to that ORF compared with all other cells at Day 0 and Day 18. Each point represents one ORF, with cells grouped by all barcodes associated with that ORF. The *x*-axis shows the mean log1p-normalised expression of each ORF in cells not assigned to that exogenous ORF, and the *y*-axis shows the mean log1p-normalised expression in cells assigned to one of the corresponding ORF-linked barcodes. Shaded contours indicate point density. The dashed diagonal line indicates equal expression between ORF-assigned and other cells. ORFs above the diagonal are more highly expressed in cells assigned to their corresponding ORF-linked barcodes. **e,** Heatmaps comparing barcode-based ORF assignment with expression of the corresponding ORFs at Day 0 and Day 18. Rows indicate ORF calls assigned by barcode identity, and columns indicate TF gene expression. Expression values for each ORF were scaled to a maximum of 1 per column. Only ORFs assigned to at least 10 cells were included, retaining 360 ORFs at Day 0 and 549 ORFs at Day 18. For clarity, only a subset of transcription factors is labelled because of space limitations. The underlying data for the Day 0 and Day 18 heatmaps are provided in Supplementary Table 5 and 6, respectively. **f,** Histogram showing the rank percentile of the diagonal value for each ORF-call row in the heatmaps shown in **e** at Day 0 and Day 18. For each row, the matched ORF expression value on the diagonal was ranked against all ORF expression values in that row. Lower percentile values indicate stronger concordance between barcode-based ORF assignment and expression of the corresponding ORF.

#### Extended Data Fig. 7

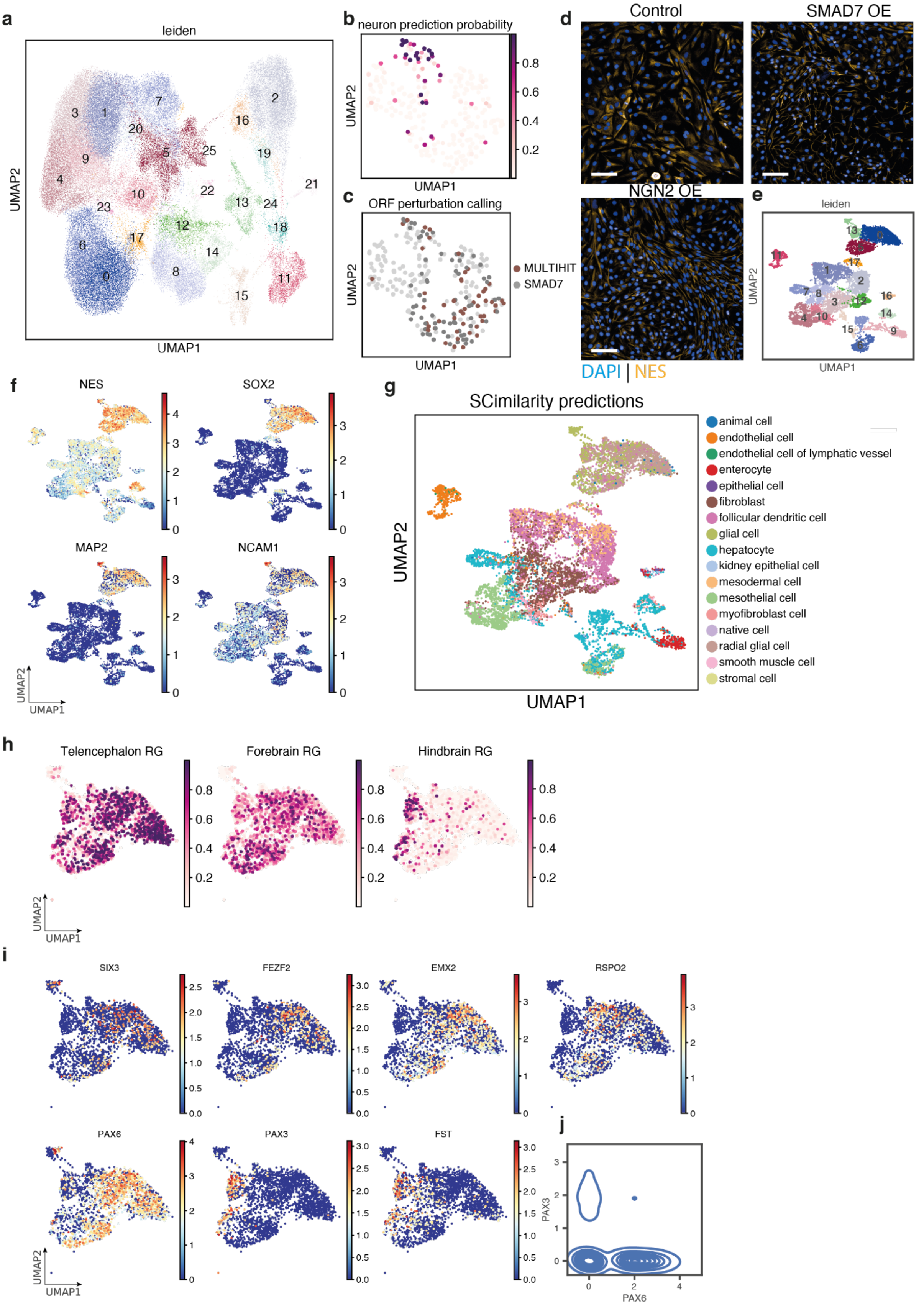

**Extended Data Fig. 7 | *SMAD7* induces neuronal differentiation.**

**a,** UMAP embedding of all D18 cells from the pooled screen, coloured by Leiden cluster (same embedding as shown in Fig. 5d, left). **b and c,** UMAP embeddings of cells from Leiden cluster 22 (same embedding as shown in Fig. 5d right), coloured by CellTypist prediction probability for neurons using the Pan Fetal Human model[43] in **b**, or by *SMAD7* or MULTIHIT barcode assignment in **c**. Cluster 22 contained 70 cells assigned to *SMAD7*-associated barcodes and 49 cells assigned to MULTIHIT barcodes, where MULTIHIT denotes barcodes linked to more than one transgene in the PacBio library. Of the MULTIHIT cells, 27 were assigned to MULTIHIT-GGTCTCCATACG-602, for which the barcode GGTCTCCATACG was linked predominantly to *SMAD7* in the library (480 reads for *SMAD7* and 122 for *BCL11A*). The remaining 22 MULTIHIT cells were assigned to MULTIHIT-ACCTGCCGCTTC-2209, for which the barcode ACCTGCCGCTTC was also linked predominantly to *SMAD7* (2,194 reads for *SMAD7* and 15 for TGIF2LX). These results suggest that the MULTIHIT cells in cluster 22 were most likely also derived from *SMAD7* overexpression. **d,** Immunostaining images of wild-type control, SMAD7- and NGN2-overexpressing cells harvested at Day 18 stained for DAPI and NES (scale bar: 100 μm). **e,** UMAP embedding of cells harvested at D18 from wild-type and *SMAD7*-overexpression cultures (same embedding as in Fig. 5f), coloured by Leiden clustering. **f,** UMAP embedding of cells harvested at D18 from wild-type and *SMAD7*-overexpression cultures (same embedding as in Fig. 5f), coloured by selected neuronal marker genes. **g,** UMAP embedding of cells harvested at D18 from wild-type and *SMAD7*-overexpression cultures (same embedding as in Fig. 5f), coloured by SCimilarity[47] prediction results. **h,** UMAP embedding of *SMAD7*-overexpressing cells (same embedding as in Fig. 5g), coloured by CellTypist prediction probabilities for each cell type using the Developing Human Brain model[48]. **i,** UMAP embedding of *SMAD7*-overexpressing cells (same embedding as in Fig. 5g), coloured by expression of selected marker genes. **j,** Kernel density plot showing the joint distribution of *PAX6* and *PAX3* expression in *SMAD7*-overexpressing cells.

#### Extended Data Fig. 8

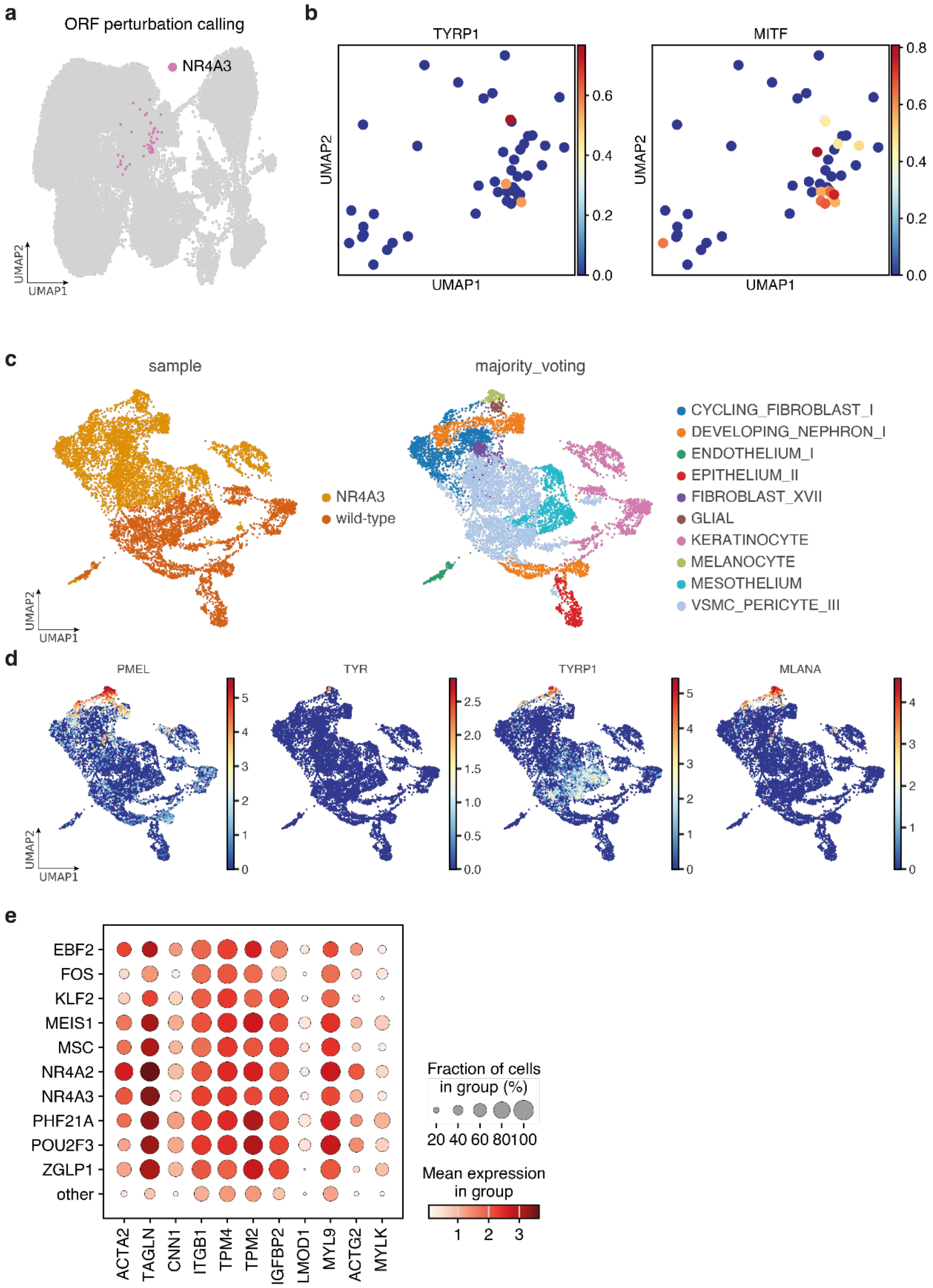

**Extended Data Fig. 8 | *NR4A3* induces melanocyte differentiation and *NR4A2* induces vSMC differentiation.**

**a,** UMAP embedding of all D18 cells from the pooled screen, highlighting cells assigned to *NR4A3*-associated barcodes. **b,** UMAP embedding of cells assigned to *NR4A3*-associated barcodes, subsetted from **a** and coloured by expression of selected melanocyte marker genes. **c,** UMAP embedding of cells harvested at D18 from wild-type and *NR4A3*-overexpression cultures, coloured by condition (left) and CellTypist majority-voting predictions using the Pan Fetal Human model[43] (right). **d,** UMAP embedding of cells harvested at D18 from wild-type and *NR4A3*-overexpression cultures (same as in **c**), coloured by expression of selected melanocyte marker genes. **e,** Dot plot showing expression of vSMC-associated marker genes across D18 cells assigned to different ORFs in Group 3, and all other D18 cells (other). Dot size indicates the percentage of cells within each ORF group expressing the gene, and dot colour indicates mean gene expression within each cell group.

#### Supplementary Table Legends

**Supplementary Table 1**
Metadata of all samples including accession numbers and metadata such as library type, cellranger versions, reference genome and sorting information.

**Supplementary Table 2**
Details of all reagents used in this study - oligonucleotides, vector and locus maps, sequences and antibodies.

**Supplementary Table 3**
Barcode mapping to the corresponding TF ORF obtained from PacBio sequencing.

**Supplementary Table 4**
Feature reference file used for analysis during Cell Ranger processing for the pooled screen to identify the ORF-barcode pairs and CITE-seq.

**Supplementary Table 5**
Normalised ORF expression matrix underlying the heatmap (Day 0) shown in Extended Data Fig. 6d, with rows corresponding to barcode-assigned ORFs and columns corresponding to expression of the matching ORFs across cells assigned to each barcode.

**Supplementary Table 6**
Normalised ORF expression matrix underlying the heatmap (Day 18) shown in Extended Data Fig. 6d, with rows corresponding to barcode-assigned ORFs and columns corresponding to expression of the matching ORFs across cells assigned to each barcode.
